# Arbuscular mycorrhiza convey significant plant carbon to a diverse hyphosphere microbial food web and mineral-associated organic matter

**DOI:** 10.1101/2023.07.11.548626

**Authors:** Anne Kakouridis, Mengting Yuan, Erin E. Nuccio, John A. Hagen, Christina A. Fossum, Madeline L. Moore, Katerina Y. Estera-Molina, Peter S. Nico, Peter K. Weber, Jennifer Pett-Ridge, Mary K. Firestone

**Author notes:** Authors for correspondence: Mary Firestone Anne Kakouridis.

## Abstract

- Arbuscular mycorrhizal fungi (AMF) transport substantial plant carbon (C) that serves as a substrate for other soil organisms, a precursor of soil organic matter (SOM), and a driver of soil microbial dynamics. Using two-chamber microcosms where an air gap isolated AMF from roots, we ^13^CO_2_-labeled *Avena barbata* for six weeks and measured. the C *Rhizophagus intraradices* transferred to SOM and hyphosphere microorganisms.
- NanoSIMS imaging, IRMS, ^13^C NMR, and SOM density fractionation showed hyphae and roots had similar ^13^C enrichment. AMF transferred 0.77 mg C per g of soil (increasing total C by 2%); 33% was found in occluded or mineral-associated pools, primarily as carbohydrates.
- In the AMF hyphosphere, there was no overall change in community diversity but 36 bacterial ASVs significantly changed in relative abundance. With stable isotope probing (SIP)-enabled shotgun sequencing, we found taxa from the Solibacterales, Sphingobacteriales, Myxococcales and Nitrososphaerales (ammonium oxidizing archaea) were highly enriched in AMF-imported ^13^C (>20 atom%). Mapping ^13^C-enriched metagenome-assembled genomes to total ASVs showed at least 92 bacteria and archaea were significantly ^13^C-enriched.
- Our results illustrate the quantitative impact of hyphal C transport on the formation of potentially protective SOM pools and indicate microbial roles in the AMF hyphosphere soil food web.

## Introduction

Arbuscular mycorrhizal fungi (AMF, phylum Mucoromycota, subphylum Glomeromycotina) form symbiotic associations with over 80% of vascular plant families (Schüβler *et al*., 2001; Spatafora *et al*., 2016) and facilitate plant nutrient uptake in exchange for photosynthetically derived carbon (C) (Smith & Read, 2008; Willis *et al*., 2013). AMF consume up to 20% of plant photosynthate C and grow extensive hyphal networks into soil (Jakobsen & Rosendahl, 1990; Tome *et al*., 2015), which can account for 15-30% of the soil microbial biomass (Leake *et al*., 2004; Parniske, 2008; Qin *et al*., 2017), and explore a soil volume substantially larger than fine roots alone (See *et al*., 2022). The composition, exudates, and interactions of AMF hyphae with plant roots and soil microbes play a critical and complex role in soil C processes (Wei *et al*., 2019; Domeignoz-Horta *et al*., 2021; Horsch *et al*., 2023). While soil C dynamics have been the focus of substantial research, gaps remain in our knowledge of the processes and factors that govern the magnitude and fate of C fluxes within the mycorrhizal pathway (Sulman *et al*., 2018; Domeignoz-Horta *et al*., 2021) and how surrounding soil biota might benefit.

Plant-fixed C, distributed into soil as root biomass and exudates, is the primary source of soil organic C and is transformed into soil organic matter (SOM) through diverse chemical and microbial processes (Torn *et al*., 1997; Trumbore, 2000; Schmidt *et al*., 2011). Since SOM holds not only organic C but also water and nutrients, its persistence is a major goal in climate change mitigation and sustainable land management. The mechanisms responsible for SOM persistence are complex and under active investigation (Jastrow *et al*., 2007; Treseder, 2016; Lehmann *et al*., 2020; Dynarski *et al.,* 2020). Amidst the paths between plant photosynthate and SOM, AMF hyphae and soil microorganisms serve as primary intermediaries; their biomass and residues can contribute to slow-cycling and persistent forms of C that may become associated with mineral surfaces or occluded within aggregates (Miller & Jastrow, 2000; Dynarski *et al.,* 2020; Angst *et al.,* 2021; See *et al*., 2022). While occluded and mineral-associated soil C is not always more persistent (Keiluweit *et al.,* 2015; Li *et al.,* 2021), mineral-associated C does typically have longer turnover times than other soil pools (Heckman *et al.,* 2022).

Several methods can distinguish C present as mineral-associated or aggregate-occluded C-forms. In the ‘density fraction’ approach, SOM is partitioned based on density into soil fractions that are operationally defined: particulate organic matter (free light fraction, FLF), occluded within soil aggregates (occluded light fraction, OLF), and mineral-associated (heavy fraction, HF) (Sollins *et al*., 2006, 2009). Organic C in these soil fractions has distinct rates of biochemical and microbial degradation (Sollins *et al*., 2006, 2009), and these fractions are widely thought to represent ecologically relevant soil components (Moni *et al*., 2010; Hatton *et al*., 2012) shaped by climate and ecosystem type (Sokol *et al.,* 2022).

AMF may alter the relative distribution of organic C in different soil fractions (Orwin *et al*., 2011; Cheeke *et al*., 2016; Frey, 2019; Soudzilovskaia *et al*., 2015, 2019) and thereby influence the persistence of C in soil. AMF have been shown to increase (Treseder, 2016; Wang *et al*., 2016; Zhang *et al*., 2020) or decrease (Hodge *et al*., 2001; Cheng *et al*., 2012; Herman *et al*., 2012; Paterson *et al*., 2016; Frey 2019) soil C accumulation depending on the study system, environmental variables, and time frame. AMF respire C and can lead to an increase in decomposition and soil respiration (Hodge *et al*., 2001; Cheng *et al*., 2012; Lang *et al*., 2021), reducing the amount of C available for SOM formation. Yet, AMF can promote soil C accumulation by increasing the formation of soil aggregates (Rillig *et al*., 2001a; Wilson *et al*., 2009; Leifheit *et al*., 2015) and SOM-mineral associations (Smits *et al*., 2009; See *et al*., 2022), and thereby protect organic C from decomposition. AMF directly contribute C to SOM formation, as their cell materials become components of SOM when they senesce and are degraded (Kögel-Knabner, 2002; Langley & Hungate, 2003; Godbold *et al*., 2006). AMF hyphae also release C compounds into surrounding soil (hyphosphere) as exudates (Toljander *et al*., 2007; Hooker *et al*., 2007) that can be transferred to other microbes (Herman *et al*., 2012; Kaiser *et al*., 2015; Bunn *et al*., 2019).

The presence of AMF may quantitatively and qualitatively alter soil bacterial communities and AMF hyphae and spores provide important niches for bacterial interactions and growth. Bacteria can colonize vital hyphae (Toljander *et al*., 2006, 2007) and form biofilms on hyphal surfaces (Lecomte *et al*., 2011), consume hyphal exudates (Kaiser *et al*., 2015), and help AMF mobilize nutrients in soil (Jiang *et al*., 2021). Bacteria can also attach to non-vital hyphae and use them as a substrate (Toljander *et al*., 2006, 2007). Our prior work suggests AMF have diverse effects on nearby hyphosphere microbiomes: stimulating organic matter decomposition and nitrogen (N) transfer (Nuccio *et al*., 2013), supporting water transport (Kakouridis *et al*., 2022) and drought resilience (Hestrin *et al*., 2022), and stimulating cross-kingdom trophic interactions (Nuccio *et al*., 2022). However, these small-scale interactions can be difficult to directly measure, and under native soil conditions, it is particularly challenging to distinguish root/rhizosphere effects from hyphosphere effects.

AMF represent a globally important pathway for the flow of C from plants into soil (Hawkins *et al*., 2023), yet it remains unclear how AMF and their hyphosphere microbiome influence the initial incorporation of organic C into soil, separate from roots. In this study, we used ^13^C stable isotope tracing and molecular techniques to measure C transfers into soil, from the host plant *Avena barbata*, a widespread annual grass, via the AMF *Rhizophagus intraradices*. In a greenhouse experiment, we used a two-chamber microcosm design to isolate root transfer of C from AMF transfer into soil. To assess how AMF affect the short-term fate of plant-derived C, we tracked ^13^C after it was fixed by host plants and transferred by AMF during six weeks of exponential phase plant growth. We characterized the form of AMF-contributed C by density fractionation into FLF (likely still free hyphae), OLF (contained within aggregate structures), and HF (mineral-associated) pools. Changes in C chemistry were assessed by ^13^C nuclear magnetic resonance (NMR) spectroscopy of soil aggregates. We also investigated the influence of AMF on hyphosphere soil microbial communities (independent from roots) by amplicon sequencing soil DNA and identifying the key microbial consumers of hyphal-derived C via stable isotope probing (SIP)-enabled metagenomic sequencing and genome reconstruction.

## Materials and Methods

### Experimental Set-up

Our experimental set-up is described in detail in Kakouridis *et al*. (2022). In brief, three two-week old *A. barbata* seedlings were planted in the ‘plant compartment’ of two-compartment microcosms (10×2.5×26.5 cm; Fig. **1a**) which was separated from the ‘no-plant compartment’ by a 3.2 mm air gap to prevent liquid water from travelling passively between compartments. Both sides of the air gap had nylon mesh, either 18 μm (allowing hyphae but excluding roots), or 0.45 μm (excluding both hyphae and roots). A total of twenty-four microcosms were used, eighteen with 18 μm mesh and six with 0.45 μm mesh. The plant compartment was packed with a 1:1 sand-clay mixture to a 1.21 g/cm^3^ density (referred to herein as the ‘sand mix’). The no-plant compartment (10×1×26.5 cm) was packed with a 1:1 soil-sand mixture to a 1.21 g/cm^3^ density (referred to as the ‘soil mix’). Soil (0 to 10 cm) was collected at the Hopland Research and Extension Center (38°59′35″N, 123°4′3″W) where *A. barbata* was the dominant vegetation.

**Figure 1:**
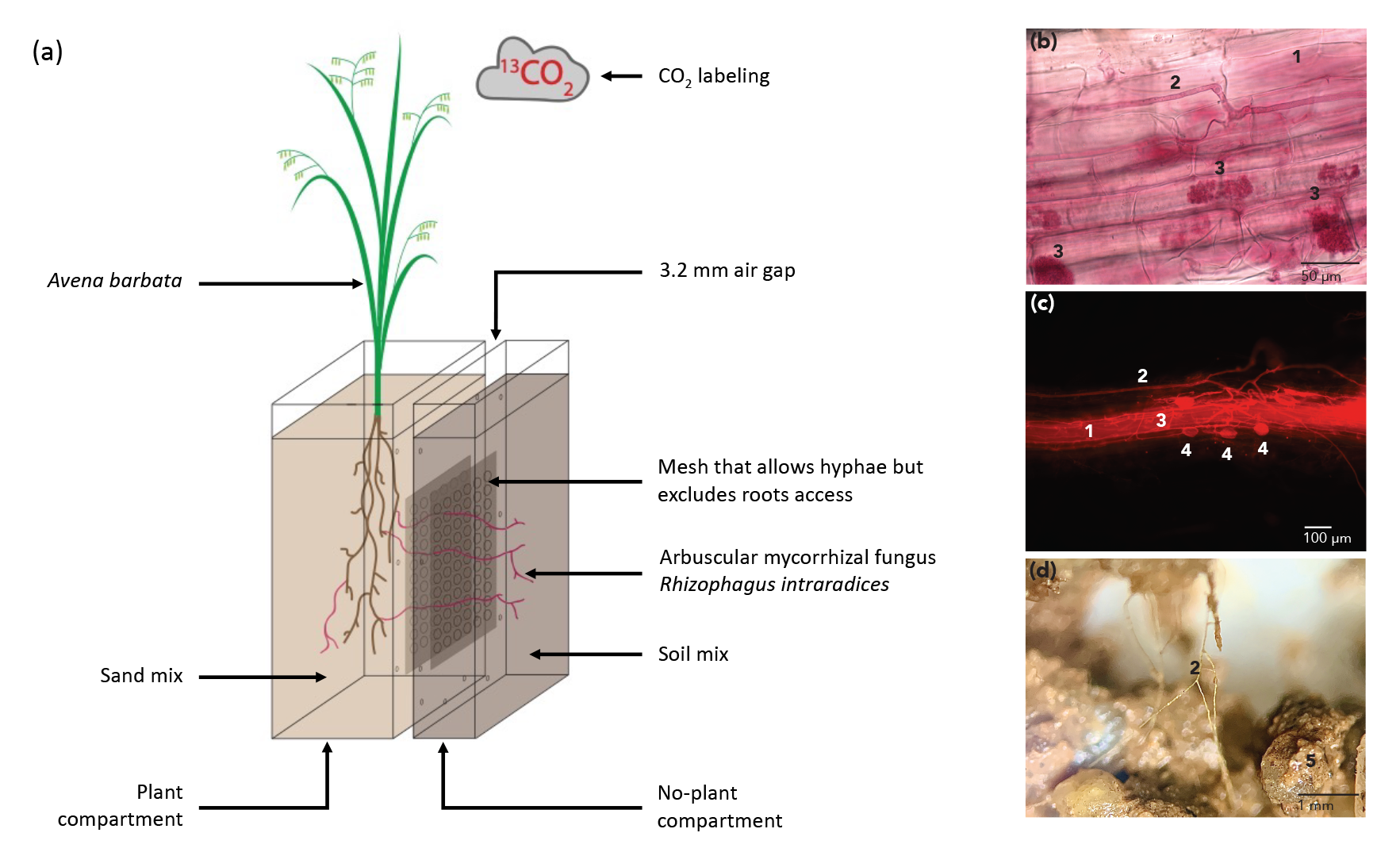
AMF, inoculated within the plant compartment, grew across an air gap to reach the no-plant compartment. **(a)** Our experimental design tested for the movement of plant C into root-free soil via AMF hyphae. This figure shows an AMF-permitted ^13^C microcosm (“+AMF”) where plants were grown in a ^13^CO_2_ atmosphere in weeks 5-10 and AMF were able to access a no-plant compartment by crossing a 3 mm air gap (through a mesh with 18-µm pore size). Two other treatments included: (1) AMF-excluded ^13^C control (“-AMF”) where plants were grown in a ^13^CO_2_ atmosphere in weeks 5-10 but AMF were not able to access the no-plant compartment because the mesh (0.45-µm pore size) between the compartments restricted both roots and hyphae from crossing from the plant compartment, and (2) AMF-permitted ^12^C control (“^12^C”), where plants were grown with an ambient CO_2_ atmosphere in weeks 5-10 and AMF were able to access the no-plant compartment. All microcosms experienced ambient CO_2_ atmosphere in weeks 1-4. Plants were harvested at the end of week 10. In sum, the experiment included 12 +AMF microcosms, 6 -AMF microcosms, and 6 ^12^C microcosms. **(b-d)** *Avena barbata* roots stained with acid fuchsin showing AMF structures. **(b)** Bright field micrographs. **(c)** Fluorescence images at wavelengths λ_ex_ 596 nm and λ_em_ 615 nm, which target AMF. **(d)** Soil-sand mixture from the no-plant compartment of +AMF microcosms with AMF hyphae visible under a dissecting microscope. In (b-d): **1**, Root; **2**, Hypha; **3**, Arbuscule; **4**, Vesicle.

In the plant compartment, the sand mix was inoculated with 26 g of *R. intraradices* whole inoculum (accession number AZ243, International Culture Collection of (Vesicular) Arbuscular Mycorrhizal Fungi (INVAM), The University of Kansas, Lawrence, KS). Bone meal (78 mg) was mixed into the plant compartment to encourage AMF establishment, and into the no-plant compartment to act as a bait for AMF to cross the air gap (Table **S1**).

The microcosms were incubated in growth chambers in the Environmental Plant Isotope Chamber (EPIC) facility, located in the Oxford Tract Greenhouse at UC Berkeley, where environmental conditions were monitored and controlled. Three chambers were used, with eight microcosms in each, organized in a randomized fashion. Volumetric water content was monitored with electronic probes (EC-5, Decagon Services, Pullman, WA, USA), and maintained at approximately 17% by watering three times weekly with autoclaved distilled water in both compartments. 10 mL of filter-sterilized Rorison’s nutrient solution (Rorison & Rorison, 1987) was added to the plant compartment (low P) and no-plant compartment (high P) once per week (Table **S1**). The plant compartment of microcosms with 0.45 μm mesh received twice as much nutrient solution as microcosms with 18 μm mesh, to make up for the nutrients plants could obtain via AMF from the no-plant compartment (Table **S1**).

Twelve microcosms with 18 μm mesh (AMF-permitted ^13^C microcosms, termed ‘+AMF’) and six microcosms with 0.45 μm mesh (AMF-excluded ^13^C microcosms, termed ‘- AMF’) were placed in a ^13^C-labeled CO_2_ atmosphere during weeks 5-10. The remaining six microcosms with 18 μm mesh (AMF-permitted ^12^C microcosms, termed ‘^12^C’), remained in a natural abundance CO_2_ atmosphere for the full ten weeks.

### Harvest and Sample Processing

At week 11, all microcosms were destructively sampled. Shoots were cut at the base, dried at 60°C and weighed for above ground biomass (Table **S1**). Roots were gently harvested and divided into three aliquots, so that each aliquot contained a randomized subsample of roots representing one third of the root system: (1) roots for staining with acid fuchsin were placed in distilled water, (2) roots for molecular analysis were placed in cell release buffer (Brodie *et al*., 2011), and (3) roots for ^13^C IRMS analysis and roots for below ground biomass measurements were placed in paper envelopes and dried 60°C (Table **S1**).

The sand mix and soil mix were collected and split into several aliquots: (1) 10 g of for gravimetric water content was oven dried at 105°C (Table **S1**) and (2) samples for molecular analyses and nutrient measurements were flash frozen in liquid nitrogen and stored at -80°C. In addition, soil mix was preserved for hyphal extraction by storing at 4°C and for soil density fractionation by air drying.

Hyphae evident in the air gap were collected on the mesh facing the inside of the air gap using tweezers and placed into tubes for DNA extraction the same day. Visible hyphae were also collected from the plant and no-plant compartments of each microcosm using a dissecting scope and tweezers, then placed on silica wafers covered with carbon sticky tape for scanning electron microscope (SEM) and nanoscale secondary ion mass spectrometry (NanoSIMS).

### Microscopy

Roots were stained with acid fuchsin using a protocol modified from Habte & Osorio (2001) and described in Kakouridis *et al*. (2022). Stained roots were mounted on slides and observed under both bright field and fluorescence (λ_ex_ 596 nm/ λ_em_ 615 nm).

### Molecular Methods

To extract AMF spores and hyphae from the sand and soil mix, and then extract DNA from roots, soil mix, spores and hyphae, we followed methods described in Kakouridis *et al*. (2022). In brief, we used a cell release buffer (Brodie *et al*., 2011) to remove surface microbial cells from roots, hexametaphosphate to separate out AMF spores and hyphae, and conducted DNA extraction from roots, spores, hyphae, and soil mix with a DNeasy PowerSoil kit (Qiagen). To collect hyphosphere microbial communities, hyphae with soil attached were picked using tweezers under a dissecting microscope.

DNA extracted from roots, soil mix, spores and hyphae, and hyphosphere microbial communities was quantified with the Quant-iT™ PicoGreen™ dsDNA Assay Kit (Invitrogen) and concentrations were normalized to 5 ng/μL. To identify the AMF present, PCR was conducted using WANDA (Dumbrell *et al*., 2011) and AML2 (Lee *et al*., 2008) primers according to procedures described in Kakouridis *et al*. (2022). A sequence was considered a match for *R. intraradices* if query coverage and percent identity were both greater than 97%.

DNA extracts were used for both amplicon and SIP-metagenome sequencing. To identify the bacteria present in hyphae and hyphosphere soil, a sequencing library was prepared using a phasing amplicon technique (Wu *et al*., 2015) on the normalized DNA samples from +AMF and -AMF soil mix from the no-plant compartment with 515F/806R primers (Caporaso *et al*., 2012) targeting the 16S rRNA gene V4 region. The library was prepared in the Zhou laboratory at the University of Oklahoma and sequenced on the Illumina MiSeq platform with 2×250bp paired-end format.

### 16S rRNA Sequence Processing

Amplicon sequences were processed with Qiime2 (Bolyen *et al*., 2019). After demultiplexing, a total of 637,832 raw sequences were obtained from the six +AMF and six - AMF soil samples. After primer trimming, sequences were denoised using DADA2 (Callahan *et al*., 2016) and clustered into amplicon sequence variants (ASVs). The representative sequences of each ASV were then used to assign taxonomy based on a classifier trained with the SILVA database (version 132-99-515-806). ASVs unassigned at the Domain level or identified as mitochondria or chloroplast sequences were discarded. This resulted in 3,019 ASVs in 12 samples, with 23,502 to 41,551 sequences per sample.

### 13C DNA-SIP and Metagenomic Read Processing

_13_C DNA-SIP, metagenomic sequencing, and metagenome assembled genome (MAG) assembly of the fractionated DNA samples have been described previously by Nuccio *et al*. (2022). Briefly, 350 ng of ^13^C-and ^12^C-AMF DNA (n = 3) were added to SIP density gradients, ultracentrifuged, fractionated, precipitated, and quantified with Lawrence Livermore National Laboratory’s high-throughput ‘HT-SIP’ pipeline (Nuccio *et al*., 2022).

After quality control, we retrieved 16S sequences from the SIP-metagenomics dataset by mapping the raw SIP-metagenome reads (14 fractions per gradient) to our 3,019 16S amplicon ASVs using bbsplit (Bushnell, 2014); this strategy allowed us to identify ^13^C-enriched organisms that may not have assembled in our previous work (Nuccio *et al*., 2022). Sequences were required to unambiguously map to a single ASV and mapped in “semiperfect” mode, which requires a 100% sequence match (max index of 2) but allows N’s and will allow a read to run off the end of the sequence. We generated an OTU table from the bbsplit ASV matches for each SIP fraction, and then calculated ASV atom percent excess (APE) using the ‘quantitative SIP’ (qSIP) pipeline (Hungate *et al*., 2015; Koch *et al.,* 2018). We required an ASV be present in 3 replicates and at least 3 fractions. To determine if an ASV was significantly enriched in ^13^C, we required the mean APE lower 90% confidence interval be significantly greater than zero, and the weighted average differences (WAD) of the ^13^C and ^12^C control density curves be significantly different (p value < 0.06).

### Soil Density Fractionation

We used a soil density fractionation method described in Fossum *et al*. (2022), which was modified from Hicks Pries *et al*. (2018) and Strickland and Sollins (1987). In brief, 20 g of air-dried soil mix and 50 mL of sodium polytungstate (SPT) (Geoliquids) was prepared to a density of 1.75 g/cm^3^, centrifuged at 3,700 RCF and left to settle. Particles floating on top were defined as FLF. The FLF was aspirated onto a 0.8 μm glass microfiber filter (Wattman), rinsed with Milli-Q water and then dried at 55 °C and weighed. To collect the OLF, the remaining soil-SPT mixture was shaken with a benchtop mixer for 1 minute, sonicated for 90 seconds, allowed to settle, and then centrifuged for one hour. The OLF was then removed by aspiration, dried, and weighed. The remaining sediment (ρ > 1.75 g/cm^3^) was defined as the HF. 150 mL of Milli-Q water was added, vigorously shaken by hand, then centrifuged for 20 minutes. The supernatant was aspirated and discarded. This process was repeated 5 times, or until the density of the supernatant reached 1 g/cm^3^. The HF was transferred, dried and weighed in the same manner as the FLF and OLF. After drying, all fractions were ground to a fine powder.

### 13C isotope ratio mass spectrometry (IRMS)

Samples of shoots, roots, sand mix, soil mix, and soil fractions were finely ground, weighed, and analyzed for total C and ^13^C abundance by dry combustion on a PDZ Europa ANCA-GSL elemental analyzer interfaced to a PDZ Europa 20-20 isotope ratio mass spectrometer (Sercon Ltd., Cheshire, UK) with a precision for ^13^C measurements of 0.1 per mil.

### Nano-scale secondary ion mass spectrometry (NanoSIMS)

To measure the atom% ^13^C of hyphae, mounted hyphae collected from the plant compartment, air gap, and no-plant compartment were coated with gold and mapped with an SEM (Inspect F, FEI, Hillsboro, OR). The isotopic composition of the hyphae was measured by NanoSIMS (NanoSIMS 50, CAMECA, Gennevilliers, France). We collected a total of 150 analysis regions (30 μm by 30 μm) to gain a broad survey of the distribution of ^13^C enrichment in hyphae. Each location was first sputtered with a ∼60 pA Cs^+^ beam to a depth of approximately 60 nm to enhance the yield of negative secondary ions and reach sputtering equilibrium (stable secondary ion counts; Ghosal *et al*., 2008). Then a 2 pA Cs^+^ beam with a nominal spot size of 200 nm was used to raster the analysis area with 256 x 256 pixels with 20 – 30 scans per sample. Five masses were collected: ^16^O^−^, ^12^C_2−_, ^12^C^13^C^−^,^12^C^14^N^−^, and ^31^P^−^using electron multipliers and a mass resolving power ∼7,000 (1.5× correction; Pett-Ridge & Weber 2012). *Pseudomonas stutzerii* cells with known isotopic composition (previously analyzed by IRMS) were used as standards.

Images were processed using L’Image software (L Nittler, www.limagesoftware.net). Images were corrected for deadtime and drift. Regions of interest (ROIs) were selected by first identifying hyphal structures from SEM and secondary electron imaging, and then based on areas of relatively uniform ^13^C enrichment from ratio images of ^12^C^13^C^−^:^12^C_2−_. Areas of high O^-^ counts (marker for minerals) were avoided. A total of 150 images were taken, from which a subset of 37 images with little charging and low O^-^ counts were kept for calculations of hyphae atom% ^13^C.

### Solid state ^13^C NMR spectroscopy

For +AMF microcosms and -AMF microcosms, hand-picked dried soil aggregate samples were finely ground and then pooled two by two as follows: samples for microcosms 1 and 2 together, 3 and 4 together, 5 and 6 together, 7 and 8 together, 9 and 10 together, and 11 and 12 together. 100 mg of each of the six pooled samples were analyzed on a 500 MHz Bruker Avance 1 Spectometer at the UC Davis NMR Facility as follows: MAS spinning speed 12 kHz, _13_C-^1^H contact time 2 ms, relaxation delay time 1 second, number of scan 60,000 to 90,000 (signal-to-noise ratio: 15∼20).

### Statistical Analyses

Statistical analyses were conducted using R version 3.6.1. (R. Core Team. 2017). A one-way analysis of variance (ANOVA) coupled with Fisher’s least-significant difference (LSD) test (package *agricolae*, p adjustment using “Holm”) was used to differentiate means of atom% and the amount of ^13^C from different treatments.

A paired Wilcoxon test was used to compare the average atom% ^13^C of roots (IRMS) versus the average of hyphae NanoSIMS measurements from +AMF microcosms (Table **S3)**. Microcosm 5 was excluded due to a technical issue with the hyphal sample preparation.

We calculated the alpha diversity (richness, Shannon index, and evenness, package *vegan*) and Faith’s phylogenetic diversity (package *picante*) of 16S rRNA detected prokaryotic communities (Oksanen *et al.,* 2019; Kembel *et al.,* 2010), and tested their differences between +AMF and -AMF treatments using a two-way ANOVA, treating “chamber” as a random factor (and no significant chamber effect was observed). P-values were corrected using false discovery rate. The principal component analysis (package *ape*) and permutational multivariate analysis of variance (Adonis based on Bray-Curtis distance, package *vegan*) were performed to evaluate community composition difference among treatments. DESeq analysis (Love *et al*., 2014) was used to identify ASVs responding to the presence of AMF. To reduce potential noise in modeling fitting, we removed ASVs that only occurred in one of the six biological replicate samples under either +AMF or -AMF conditions. After this prevalence filtering, 1,534 ASVs were included in the DESeq analysis for parameter estimation and model fitting.

## Results

### Confirmation of root colonization and hyphal networks in the no-plant compartment

Using WANDA-AML2 amplicon sequencing, we confirmed that roots of all microcosms were colonized by *R. intraradices* and confirmed this AMF in air gap hyphae and soil mix samples from +AMF and ^12^C microcosms. We also observed hyphae, spores, and arbuscules in roots stained with acid fuchsin using light and fluorescence microscopy (Fig. **1b-c**), confirming that *R. intraradices* was actively growing in roots over the course of our experiment. We observed hyphae crossing the air gap, and extensive hyphal networks in the no-plant compartment of +AMF and ^12^C microcosms (Fig. **1d**). In -AMF microcosms, we did not observe hyphae crossing the air gap nor hyphal networks in the soil mix.

### Confirmation of ^13^C labeling and ^13^C presence in no-plant compartment of +AMF microcosms

The atom% ^13^C of shoots in +AMF and -AMF microcosms was 43.5 ± 1.5% and 43.7 ± 1.7%, respectively, and 1.1 ± 0.001% in ^12^C microcosms (Fig. **S1**, Table **S1**). Root enrichment was 41.3 ± 1.9 atom% and 42.2 ± 2.0 atom% in +AMF and -AMF microcosms, respectively, and 1.1 ± 0.002% in ^12^C microcosms (Fig. **S1**, Table **S1**). The atom% ^13^C of the sand mix (plant compartment) was 5.3 ± 0.4% and 4.2 ± 0.2% in +AMF and -AMF microcosms, respectively, and 1.1 ± 0.003% in ^12^C microcosms (Fig. **S1**, Table **S1**). The soil mix (no-plant compartment) of +AMF microcosms was significantly more ^13^C enriched than that of -AMF and ^12^C microcosms (atom% ^13^C 1.8 ± 0.1% versus 1.1 ± 0.001% and 1.1 ± 0.0003%, respectively, *P* < 0.001) (Fig. **S1**, Table **S1**). These results confirm there was significantly more ^13^C in the no-plant compartment of +AMF microcosms and that the no-plant compartments of -AMF microcosms were not contaminated by ^13^C. The presence of AMF did not change soil gravimetric water content, aboveground biomass, root:shoot ratio, shoot C:N ratio and %N, but significantly increased plant P content and decreased plant N:P ratio (Table **S1**; Kakouridis *et al.,*2022).

### 13C was found in light, occluded, and heavy fractions of SOM

SOM from the soil mix (no-plant compartment) was separated into FLF, OLF and HF using a sodium polytungstate density gradient approach. Based on ^13^C IRMS analysis of these density fractions, after six weeks of labeling, 26.7±3.0 mg of ^13^C was present in the no-plant compartment soil mix of +AMF microcosms (Fig. **2a**, Tables **S1 and S2**). This represents 74.4 mg C which is equivalent to 0.77 mg C per g of the native soil material (not including the sand) or 2.05% of the total C in the soil mix in the no-plant compartment (Methods **S1**).

**Figure 2:**
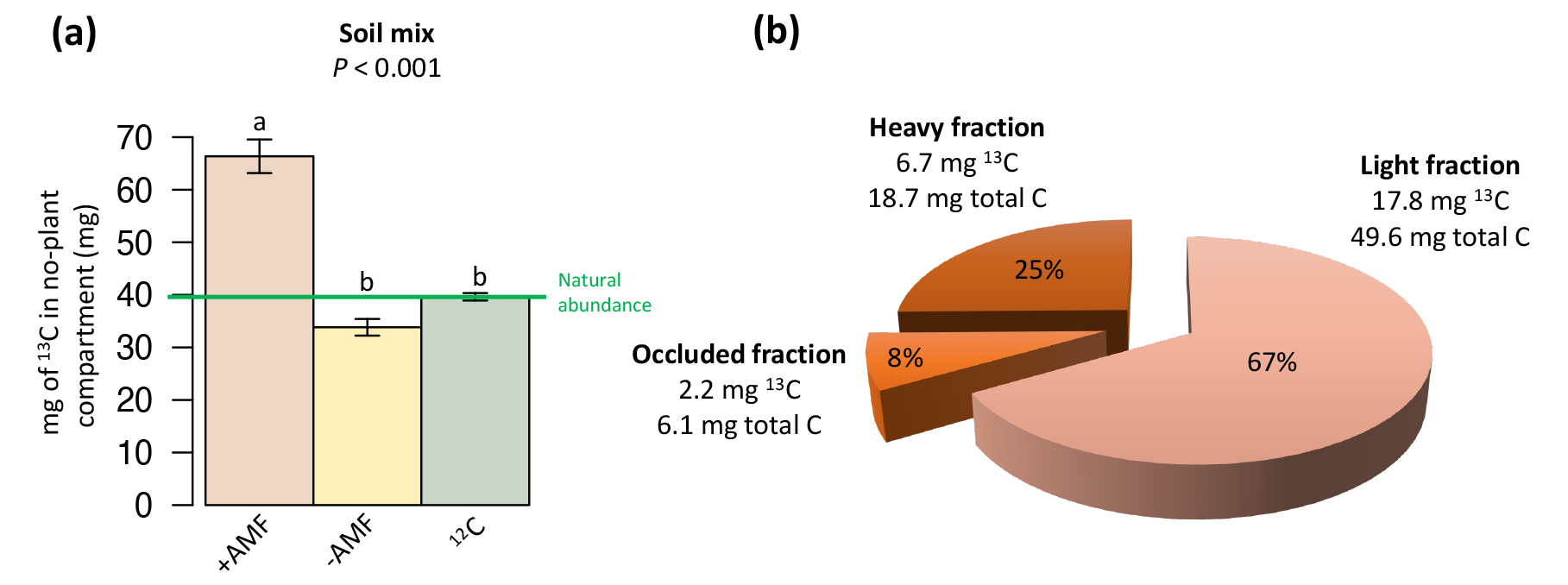
^13^C and total C transported by AMF hyphae into a no-plant compartment, by soil organic matter fraction. **(a)** mg ^13^C in soil mix (no-plant compartment) in +AMF, -AMF, and ^12^C microcosms after six weeks of labeling. Different letters represent statistically significant differences; the indicated p-value is for a one-way ANOVA & Fisher LSD test. Green line labeled “N” represents natural abundance ^13^C. **(b)** Distribution of ^13^C contributed by AMF hyphae in light, occluded, and heavy soil fractions after six weeks of labeling based on soil density fractionation followed by ^13^C IRMS analysis. Total C contributed by AMF hyphae to these soil density fractions were estimated based on the ^13^C enrichment of AMF hyphae (see Table S2 and Method S1 for calculations).

The 26.7 mg of newly added ^13^C found in the soil mix was distributed amongst the density fractions as follows: 17.8 mg ^13^C (representing 49.6 mg total C or 1.37% of the total soil mix C) in the FLF, 2.2 mg ^13^C (representing 6.1 mg C or 0.17% of the total soil C) in the OLF, and 6.7 mg ^13^C (representing 18.7 mg C or 0.52% of the total soil C) in the HF (Fig. **2b**).

### Hyphae enrichment

To determine the average ^13^C enrichment of hyphae, we collected NanoSIMS images of hyphae from the plant compartment, air gap, and no-plant compartment (Fig. **3**). Based on an average of 37 images, hyphae were 35.9±1.5% ^13^C enriched (Fig. **3f** and Table **S3)**. A paired Wilcoxon test between five matched roots and hyphae samples indicated no statistical difference between the enrichment of roots and hyphae (*P* = 0.063, Fig. **3f**).

**Figure 3:**
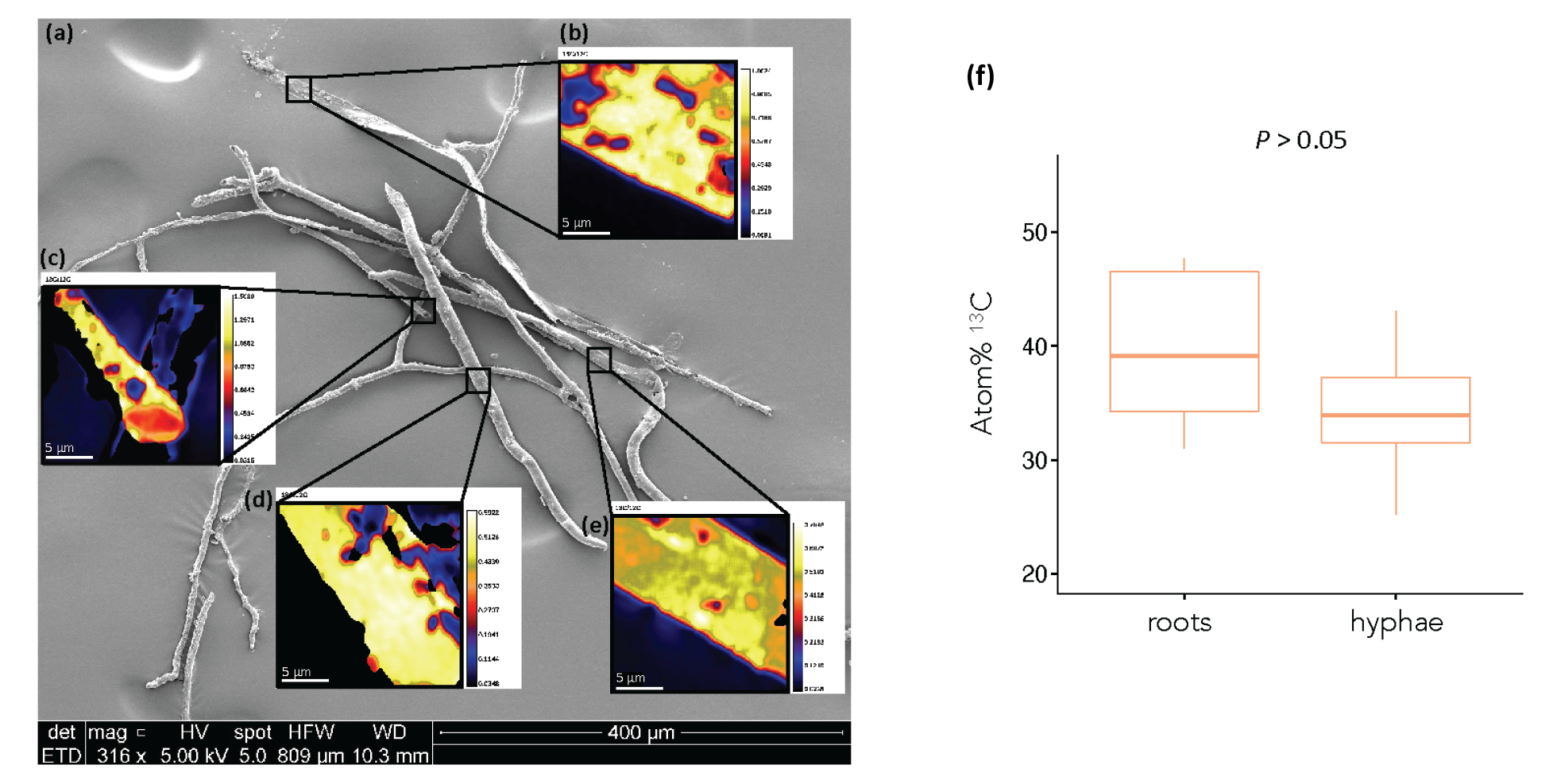
^13^C isotope enrichment of roots and hyphae. **(a)** Scanning electron microscope (SEM) and **(b-e)** nanoscale secondary ion mass spectrometry (NanoSIMS) images of hyphae from the no-plant compartment of a +AMF microcosm. **(f)** Atom% ^13^C of roots and hyphae from +AMF microcosms. Whiskers represent the minimum and maximum values in the data; *P*-value is indicated above the plot (paired Wilcoxon test, n=5).

### NMR spectra

NMR spectra were obtained for six soil mix samples (three from the +AMF microcosms and three from the -AMF microcosms) (Fig. **4**). The +AMF NMR spectra have a larger peak in the 45-110 ppm range, corresponding to the O-alkyl C functional group and indicative of carbohydrates (Kögel-Knabner, 1997) (Fig. **4** and Fig. **S2**).

**Figure 4:**
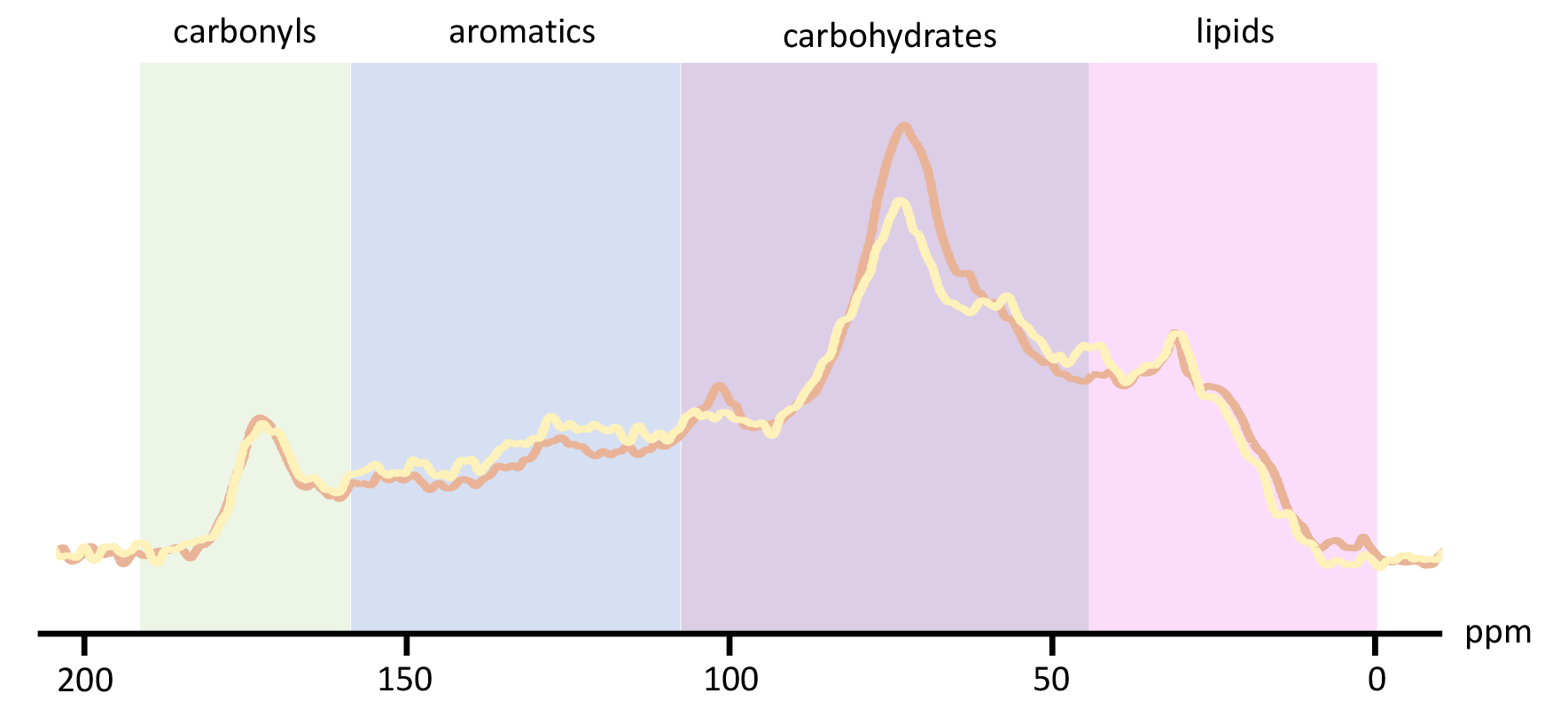
Solid state ^13^C NMR spectra of the soil mix (no-plant compartment) of a +AMF microcosm (in orange) and a -AMF microcosm (in yellow).

### Prokaryotic communities in soil mix

There were no significant differences in the number of 16S sequences obtained from +AMF (32,451 ± 4,596) versus -AMF (34,171 ± 6,655) treatments after quality control. The 12 samples analyzed resulted in 3,019 amplicon sequencing variants (ASVs), with 23,502 to 41,551 sequences per sample. There was no significant difference in the alpha diversity of prokaryotic communities in +AMF vs. -AMF microcosms, as measured by richness, Shannon index, evenness, and Faith’s phylogenetic diversity (ANOVA, *P* > 0.05). However, community composition did differ between +AMF and -AMF treatments (Fig. **5a**), measured via permutational multivariate analysis of variance (Adonis, F_1,8_ = 2.23, *P* = 0.004).

DESeq analysis identified 19 ASVs that were significantly more abundant in the presence of AMF, and 17 ASVs that were significantly less abundant in the presence of AMF (Fig. **5b** and Table **S4**). Twenty-three of these 36 (65%) ASVs belong to the Proteobacteria Phylum. Fourteen increased in relative abundance in +AMF microcosms, and nine decreased. Other ASVs that significantly increased or decreases in relative abundance in +AMF microcosms are listed in table **S4**.

### 13C enriched prokaryotes in soil mix

Of the 3,019 ASVs detected by 16S amplicon analysis, 2,934 ASVs also had a semiperfect read match within the SIP-metagenomics dataset. We identified 92 ASVs significantly enriched in ^13^C (Fig. **6**) among the total 682 ASVs that passed the qSIP criteria (e.g., detected in enough replicates and fractions). These ASVs span a diverse phylogeny, with Proteobacteria the dominant phylum. The most enriched taxa, with atom percent excess (APE) _13_C > 20%, include those from the delta-Proteobacteria order Myxococcales, Thaumarchaeota family Nitrososphaeraceae, the Acidobacteria genus Candidatus *Solibacter*, and the Bacteroidetes family AKYH767 (within the Sphingobacteriales). The most enriched ASV was from the *Haliangium* genus within the Myxococcales (APE 35.2%), an enrichment value similar to the AMF hyphae (35.9%) (note, Myxococcales has been proposed to be reorganized as the Myxococcota phylum in the GTDB taxonomy, Parks *et al*., 2018). There were 33 ASVs from diverse phylum and orders with APE of over 10% (Fig. **6**).

**Figure 5:**
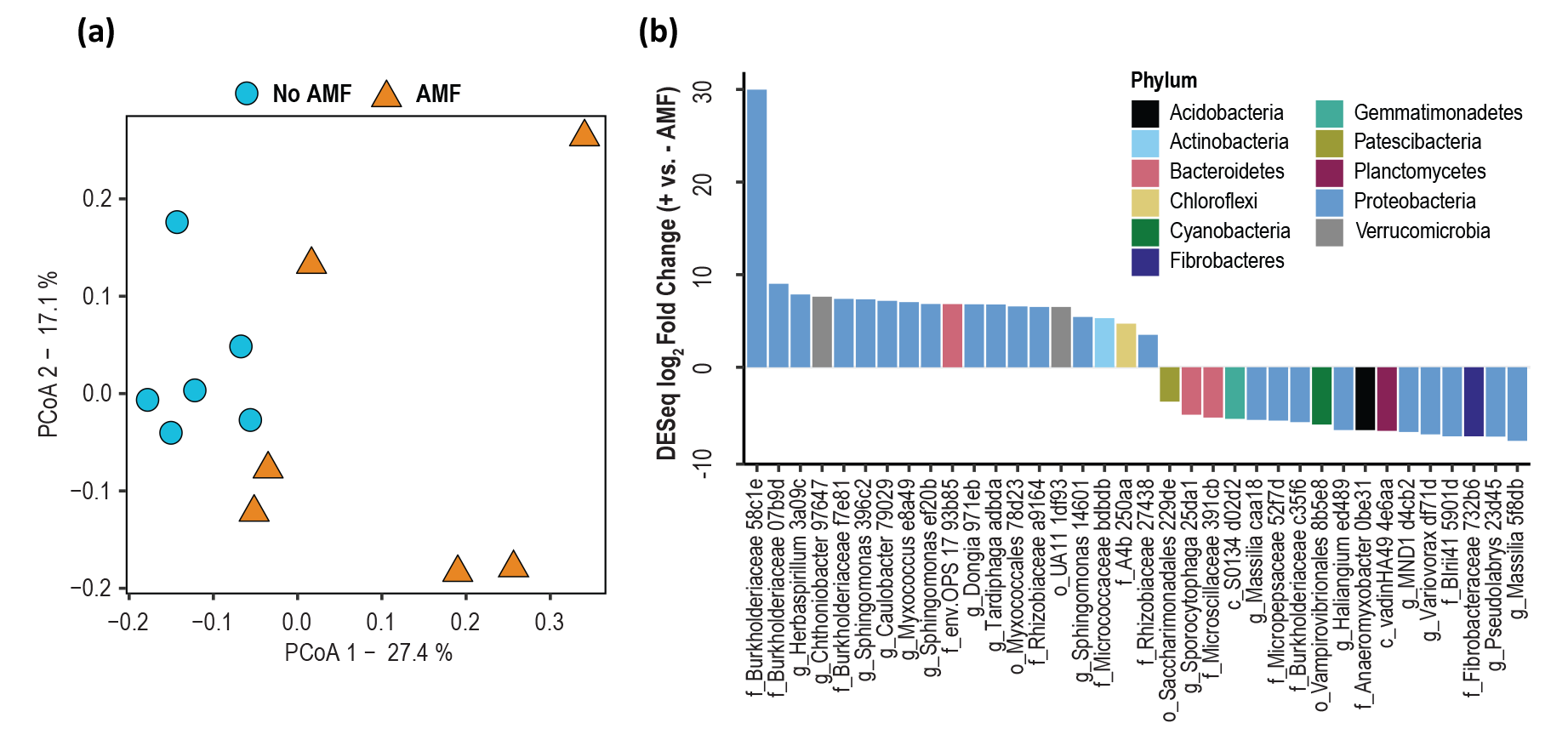
Bacterial community composition in the no-plant compartment soil mix of +AMF and -AMF microcosms based on 16S rRNA gene amplicons. (A) Principal component analysis showing distinct community composition with or without AMF. (B) Amplicon sequence variants (ASVs) that significantly increased and decreased in relative abundance +AMF microcosms compared to -AMF microcosms (DESeq analysis of 16S rRNA reads, *P* < 0.05 adjusted for multiple comparisons).

**Figure 6:**
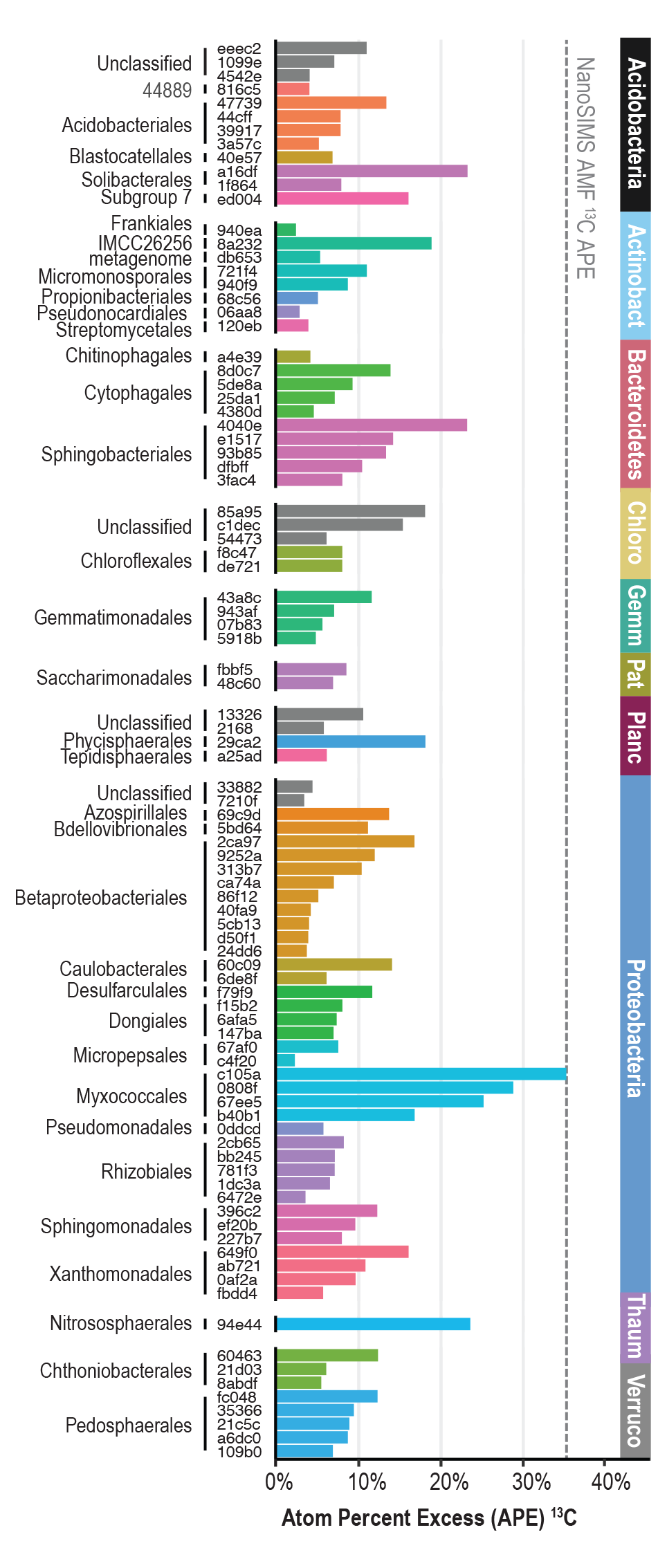
Following Stable Isotope Probing (SIP), ^13^C-enriched hyphosphere taxa and their level of enrichment in ^13^C atom% excess (APE). The ASVs (first five alphanumeric characters shown) are SIP-metagenomic reads that were mapped to corresponding ASVs obtained from 16S amplicon sequencing of the same DNA samples. Bars are grouped by Phylum of the ASVs and colored by Order. The dashed line shows the APE of AMF hyphae estimated by NanoSIMS.

## Discussion

AMF transport a significant amount of photosynthetic C into the soil matrix beyond the extent of roots alone, but the contribution of AMF to soil C retention lacks experimental exploration. We used ^13^CO_2_ labeling of *Avena barbata* in a two-compartment microcosm with an air gap designed to isolate *R. intraradices* hyphae from roots and quantify the amount and form of photosynthetic C transported via AMF hyphae into soil. After six weeks of labeling, AMF hyphae crossing the airgap were ^13^C enriched at over 35 atom% and the total C transferred by hyphae and retained in soil represented over 2% of the total soil C, a third of which was found stabilized in the mineral-associated fraction or occluded within soil aggregates.

### AMF transported C quickly enters protected pools

After six weeks of labeling, the ^13^C that remained within the no-plant compartment soil included ^13^C that was fixed by plants, transferred to AMF and transported to the no-plant compartment, minus the ^13^C that left the system via soil respiration or dissolved organic carbon (DOC) in water that drained from the microcosms. 33% of this remaining ^13^C was in either the OLF or the HF. Organic C in both fractions is thought to be at least somewhat protected from decomposition by its location inside aggregates or its association with minerals (Kleber *et al*., 2007; Keiluweit *et al*., 2012; Throckmorton *et al*., 2015; Kallenbach *et al*., 2016).

The ^13^C enrichment of soil density fractions suggests that a large portion of the AMF-derived C was still present in the form of living or dead hyphae, with 67% of the ^13^C in the FLF. The density of a fungal hypha has been estimated to be 1.1 g/cm^3^ (Bakken & Olsen, 1983), so hyphae should be concentrated in the FLF unless they are in a protected form. Organic C in the FLF is readily available to microbial and physical degradation and therefore tends to have a shorter residence time in soil compared to C in OLF and HF (Kleber *et al*., 2007; Keiluweit *et al*., 2012; Throckmorton *et al*., 2015; Kallenbach *et al*., 2016).

We found that a small fraction of the ^13^C, 8%, was in the OLF inside aggregates that were sufficiently stable to retain structure and hold on to the ^13^C during the soil density fractionation process. As soil aggregation can physically protect C-rich litter from microbial degradation, an increase in aggregation may be an important mechanism for C sequestration (Rillig, 2004; Rillig & Mummey, 2006; Rillig *et al*., 2010). AMF may increase the stability of soil aggregates by producing and releasing glomalin-related soil glycoproteins (GRSP)(Wright *et al*., 1996; Rillig *et al*., 2001a, 2001b, 2002, 2010; Halvorson & Gonzalez, 2006). A recent review, however, argues that the evidence supporting the production of GRSP by AMF is primarily correlative and GRSP may have multiple non-AMF origins (Holátko *et al*., 2020). It is clear that AMF hyphae can increase soil aggregation by physically holding soil particles together in their intricate extraradical mycelium. The physical protection of C in aggregates can be visually observed in figure **1d** where soil aggregates were mechanically broken open to reveal AMF hyphae.

Lastly, about 25% of the transported ^13^C remained in soil within the HF. The origin of this mineral-associated organic C may have been hyphal exudates, bacteria that consumed the exudates, and/or hyphae that senesced and degraded to the point of being sorbed to mineral surfaces (Pett-Ridge & Firestone, 2017). The transfer of root C into fungal and bacterial cell biomass appears to be a particularly important step that precedes C stabilization on mineral surfaces (Kleber *et al*., 2007, 2015; Keiluweit *et al*., 2012; Throckmorton *et al*., 2015; Kallenbach *et al*., 2016; Pett-Ridge & Firestone 2017). In an earlier greenhouse study, we used soil collected from the same field site and the same plant genus, but used only one compartment for both roots and hyphae. We found 43% of the C from root and hyphae in the HF after 12 weeks of labeling (Pett-Ridge & Firestone, 2017). C contributed by hyphae alone was more than half of that from both roots and hyphae, despite the shorter labeling time reported here (6 weeks) compared to that of the previous study. Both sets of results demonstrate rapid interactions between root and hyphal exudates and soil minerals.

### AMF contribute a significant portion of soil labile C via biomass and exudates

In our experiment, C transported by AMF hyphae made up over 2% of the total organic C in the no-plant compartment after a six-week incubation. This amount of organic C is substantial relative to the estimate that up to 5% of the total organic C in soil is from microbial biomass (paulPaul & Clark, 1989). This comparison is necessarily approximate because we do not know the effects that the no-plant compartment characteristics (1:1 mix of soil and sand, an air gap, and only six weeks to develop) may have had relative to natural soil under equilibrium conditions. Nonetheless, this finding is consistent with estimates that AMF are 15 to 30% of the total soil microbial biomass in natural soil (Leake *et al*., 2004; Olsson & Wilhelmsson, 2000; Rillig *et al*., 2001a; Qin *et al*., 2017). AMF have been observed to spread over 11 cm away from roots in a pot culture in seven weeks (Jakobsen *et al*., 1992), which is in line with observations in our study. It appears that AMF have the ability to grow hyphal networks quickly and thus rapidly incorporate a substantial amount of C into the soil.

The ^13^C-NMR results showed a larger carbohydrate peak in the spectrum for +AMF soil compared to -AMF soil (Fig. **5**). This suggests that AMF in the soil mix away from the direct influence of roots produced hyphae and released metabolites that contained a large proportion of carbohydrates. AMF hyphae contain chitin and are known to exude diverse compounds in the hyphosphere, including sugars (in particular hexoses), amino acids, and organic acids (Toljander *et al*., 2007; Bharadwaj *et al*., 2012; Sato *et al*.; 2015; Luthfiana *et al*., 2021), so our findings are consistent with previous studies.

### Enrichment of AMF C is comparable to that of roots

The NanoSIMS results show there is no significant difference between the ^13^C enrichment of roots and hyphae (Fig. **3f**, Tables **S2** and **S3**). Hyphae have been shown not only to bring plants nutrients but also to carry up to 34% of the water transpired by plants (Ruth *et al*., 2011; Quiroga *et al*., 2019; Kakouridis *et al*., 2022). In fact, the evolution of land plants may have only been possible through the formation of mycorrhizal associations, because the algal ancestors to land plants were not adequately equipped to live on land independently (Jeffrey, 1962; Pirozynski & Malloch, 1975; van der Heijden *et al*., 2015). NanoSIMS evidence from this study further supports the idea that AMF truly act as an extension of the root system.

### Composition and activity of AMF hyphosphere microbial communities

Local-scale soil biogeochemistry is an important driver of microbial community composition (McGuire & Treseder, 2010; Wagg *et al*., 2014; Graham *et al*., 2016; Kivlin *et al*., 2020; Anthony *et al*., 2020). We were interested in how AMF affect the soil microbial community—separately from roots—which we assessed with both ^13^C-SIP analyses and amplicon 16S sequencing. These two methods ask fundamentally different questions: ^13^C-SIP asks what organisms partake in the AMF-C food web, either peripherally (low ^13^C enrichment) or centrally (high ^13^C enrichment), while amplicon sequencing asks how the presence versus absence of AMF enables dramatic (e.g., log_2_fold) changes in the relative abundance of taxa. These two methods provide critical but different perspectives on the composition and activity of hyphosphere microbial communities.

AMF frequently co-occur with bacteria in soil away from the direct influence of roots (Yuan *et al*., 2021). Here, using 16S amplicon analysis, we found the relative abundances of 19 ASVs significantly increased and 17 ASVs decreased in the presence of AMF in the root-free hyphal compartment. Over half of these changed ASVs belonged to the Proteobacteria Phylum, suggesting a potential phylogenetic coherence of bacteria whose abundances strongly respond to the presence of AMF. These 36 ASVs only comprise 1.2% of all ASVs detected, but their abundance changes were sufficiently large to shift the overall community structure, as other studies have also discovered (Nuccio *et al*., 2013; Gui *et al*., 2017; Rodríguez-Caballero *et al*., 2017; Cao *et al*., 2020; Hao *et al*., 2020).

Some of the taxa that increased in abundance did not incorporate ^13^C. Two ASVs of the Burkholderiaceae family (58c1e and 07b9d) increased the most in relative abundance among all ASVs but were not ^13^C enriched. AMF may have stimulated these organisms by mechanisms that did not involve ^13^C consumption, such as by changing local edaphic or nutrient conditions (e.g., increasing or decreasing N, P) or by non-assimilative C processes. In addition, 10 of the 19 ASVs increased in relative abundance but we were unable to determine whether they incorporated ^13^C.

Compared to amplicon analysis, ^13^C-SIP identified many more organisms in the AMF food web. DNA-SIP isolates the populations actively assimilating a labeled substrate, and does not detect DNA from inactive microbes, relic DNA fragments, or slow-growing communities. Even though only 23% of the ASVs detected via amplicon sequencing were captured by metagenomics and subsequent qSIP analysis, qSIP detected many organisms that took up ^13^C without detectable changes in relative abundance. The AMF hyphosphere is a microhabitat that can be difficult to isolate and sample without also collecting large amounts of background soil; in these instances, SIP may more sensitively identify populations responding to AMF than standard amplicon analyses. However, since shotgun metagenomic sequencing can hardly capture all 16S genes from a soil community, the true AMF food web likely contains more taxa than detected in this study.

This study goes beyond our previous work (Nuccio *et al.,* 2022) by using 16S amplicon sequences from the same DNA samples as a reference database to identify more ^13^C-enriched hyphosphere taxa within our SIP-metagenomes. Previously, we used metagenome assembled genomes (MAGs), which limits the analysis to genomes amenable to assembly. In line with our previous findings, the three most enriched ASVs, including one with ^13^C enrichment close to that of AMF hyphae, all belong to the family Myxococcales in the genus Haliangium, likely facultative predators (Nuccio *et al.,* 2022). An ammonia oxidizing archaea (AOA), Nitrososphaera was also highly enriched with 23% APE. We also identified multiple highly ^13^C enriched taxa that were not assembled into MAGs in our previous study. For example, ASVs belonging to Solibacterales, Subgroup 7, and Phycisphaerales were ^13^C enriched with over or nearly 20% APE. Solibacterales taxa are known to solubilize phosphorous (Bergkemper *et al.,* 2016), which may synergistically interact with AMF by P-C exchange (Nacoon *et al.,* 2020). Acidobacteria Subgroup 7 is reported to increase with pH (Jones *et al*., 2009) and has potential to use different carbohydrates (de Chaves *et al*., 2019). The high ^13^C enrichment of this Acidobacteria subgroup may be a combination of microhabitat change and substrate availability associated with AMF but needs further study. The Phycisphaerales ASV matched well to soil microbes from other studies (Zeglin *et al*., 2015; Addison *et al*., 2019), and our finding that it consumed AMF-derived C may be a first clue about its function in soil.

Some ASVs with moderate APE that are from Orders Rhizobiales, Caulobacterales, Sphingomonadales have been reported to associate with AMF (Agnolucci *et al*., 2018; Hoseinzade *et al*., 2016; Akyol *et al*., 2019; Hao *et al*., 2020; Yuan *et al*., 2021). Here we confirmed their consumption of hyphae-derived C. Rhizobiales and Sphingomonadales are considered rhizosphere adaptors (Lei *et al.,* 2019) and together consist of 4.8% of the total AMF hyphosphere ASVs in our study. The hyphosphere may have similar characteristics to the rhizosphere, in that low molecular weight C from hyphal exudates selectively enrich these taxa. Their moderate but not high levels of ^13^C incorporation may be limited by the relatively small portion of the cells that are spatially close to hyphae and have access to AMF C. The photosynthetic C transported by AMF thus facilitated a diverse hyphosphere microbial food web dependent on AMF hyphal C supply.

## Conclusion

Our study shows that AMF moved a substantial amount of C from plants beyond the rhizosphere. After only a few weeks, about a third of this C had made its way into mineral-associated and aggregate-occluded forms, which may have longer residence times in soil. In hyphosphere soil, we measured an increased presence of organic C compounds (such as chitin and hexoses) that are generally associated with AMF hyphae. AMF C inputs modified the bacterial community, facilitating a diverse microbial food web that incorporated hyphae derived C. This effect likely stimulated enhanced AMF (and thereby plant) access to N and P. Together, our findings indicate that AMF have the potential to be major players in climate change mitigation and sustainable land management.

## Acknowledgements

The authors thank Dr. Thomas Bruns and Dr. John Taylor for their thoughtful advice on the project, Dr. Denise Schichnes and Dr. Steven Ruzin for their guidance with microscopy, Donald J. Herman and Christina Wistrom for their assistance with the greenhouse set up, and Dr. Michael S. Page for his help with ^13^C calculations.

## Funding

This research was supported by the U.S. Department of Energy Office of Science, Office of Biological and Environmental Research Genomic Science program under Awards DE-SC0016247 and DE-SC0020163 to UC Berkeley and awards SCW1589 and SCW1678 to Lawrence Livermore National Laboratory. Work conducted at LLNL was contributed under the auspices of the US Department of Energy under Contract DE-AC52-07NA27344. A.K. was supported by the Bennett Agricultural Fellowship, the Storie Memorial Fellowship, and the Jenny Fellowship in Soil Science.

## Authors Contribution

A.K. and M.K.F. designed the experiment with the assistance of J.A.H., M.Y., K.Y.E-M, E.E.N. and J.P-R.

A.K., J.A.H, M.M, C.A.F., P.N. and P.K.W. performed the experiment with assistance from K.Y.E-M.

A.K., J.A.H, M.Y., E.E.N. C.A.F., M.M., P.N., P.K.W. and M.K.F. analyzed the data with the assistance of P.N., K.Y.E-M, and J.P-R.

A.K., M.Y., E.E.N, J.P-R and M.K.F. drafted the manuscript. A.K., M.Y., E.E.N., J.A.H., C.A.F., K.Y.E-M, P.N., P.K.W., J.P-R., and M.K.F all contributed to the final manuscript.

## Data Availability

The data that supports the findings of this study are available from the corresponding authors upon reasonable request.

## Competing Interests

The authors declare no competing interests.

## The following Supporting Information is available for this article

**Fig. S1.:** Atom% ^13^C in shoots, roots, sand mix and soil mix in +AMF, -AMF, and ^12^C microcosms.

**Fig. S2.:** Solid state NMR spectra of the soil mix (no-plant compartment) of +AMF and -AMF microcosms.

**Table S1.:** Data used in statistical analyses.

**Table S2:** Data used to calculate how much ^13^C and total C was transported by AMF in +AMF microcosms.

**Table S3:** Atom% ^13^C of hyphae from NanoSIMS measurements for the 37 samples discussed in the main text.

**Table S4.:** ASVs that significantly increased or decreased in relative abundance in the soil mix (no-plant compartment) of +AMF microcosms.

**Method S1.**: Detailed ^13^C calculations.

## Notes

### Competing Interest Statement

The authors have declared no competing interest.

## References

Addison S, Smaill S, Garrett L, Wakelin S. 2019. Effects of forest harvest and fertiliser amendment on soil biodiversity and function can persist for decades. Soil Biology and Biochemistry 135: 194–205.

Agnolucci M, Avio L, Pepe A, Turrini A, Cristani C, Bonini P, Cirino V, Colosimo F, Ruzzi M, Giovannetti M. 2019. Bacteria associated with a commercial mycorrhizal inoculum: community composition and multifunctional activity as assessed by Illumina sequencing and culture-dependent tools. Frontiers in Plant Science 9: 1956.

Akyol TY, Niwa R, Hirakawa H, Maruyama H, Sato T, Suzuki T, Fukunaga A, Sato T, Yoshida S, Tawaraya K et al. 2019. Impact of introduction of arbuscular mycorrhizal fungi on the root microbial community in agricultural fields. Microbes and Environments 34: 23–3.

Angst G, Mueller KE, Nierop KG, Simpson MJ. 2021. Plant-or microbial-derived? A review on the molecular composition of stabilized soil organic matter. Soil Biology and Biochemistry 156: 108–189.

Anthony MA, Crowther TW, Maynard DS, van den Hoogen J, Averill C. 2020. Distinct assembly processes and microbial communities constrain soil organic carbon formation. One Earth 2: 349–360.

Bakken LR, Olsen RA. 1983. Buoyant densities and dry-matter contents of microorganisms: conversion of a measured biovolume into biomass. Applied and environmental microbiology 45: 1188–1195.

Bergkemper F, Kublik S, Lang F, Krüger J, Vestergaard G, Schloter M, Schulz S. 2016 Novel oligonucleotide primers reveal a high diversity of microbes which drive phosphorous turnover in soil. Journal of Microbiological Methods 125: 91–97.

Bharadwaj DP, Alström S, Lundquist P. 2012. Interactions among *Glomus irregulare*, arbuscular mycorrhizal spore-associated bacteria, and plant pathogens under in vitro conditions. Mycorrhiza 22: 437–447.

Bolyen E, Rideout JR, Dillon MR, Bokuich NA, Abnet CC, Al-Ghalith A, Alexander H, Alm EJ, Arumugam M, Asnicar F et al. 2019. Reproducible, interactive, scalable and extensible microbiome data science using QIIME 2. Nature Biotechnology 37: 852–857.

Brodie EL, Joyner DC, Faybishenko B, Conrad ME, Rios-Velazquez C, Malave J, Martinez R, Mork B, Willett A, Koenigsberg S et al. 2011. Microbial community response to addition of polylactate compounds to stimulate hexavalent chromium reduction in groundwater. Chemosphere 85: 660–665.

Bunn RA, Simpson DT, Bullington LS, Leckberg Y, Janos DP. 2019. Revisiting the ‘direct mineral cycling’ hypothesis: arbuscular mycorrhizal fungi colonize leaf litter, but why? ISME J 13: 1891–1898.

Bushnell B. 2014. BBTools Software Package. Available from: sourceforge.net/projects/bbmap/.

Callahan BJ, McMurdie PJ, Rosen MJ, Han AW, Johnson AJ, Holmes SP. 2016. DADA2: High-resolution sample inference from Illumina amplicon data. Nature methods 13: 581–583.

Cao J, Feng Y, Lin X, Wang J. 2020. A beneficial role of arbuscular mycorrhizal fungi in influencing the effects of silver nanoparticles on plant-microbe systems in a soil matrix. Environmental Science and Pollution Research 27: 11782–11796.

Caporaso JG, Lauber CL, Walters WA, Berg-Lyons D, Huntley J, Fierer N, Owens M, Betley J, Fraser L, Bauer M et al. 2012. Ultra-high-throughput microbial community analysis on the Illumina HiSeq and MiSeq platforms. ISME J 6: 1621–1624.

de Chaves M, Silva G, Rossetto R. Edwards R, Tsai S, Navarrete A. 2019. Acidobacteria subgroups and their metabolic potential for carbon degradation in sugarcane soil amended with vinasse and nitrogen fertilizers. Frontiers in Microbiology 10: 467603.

Cheeke TE, Phillips RP, Brzostek ER, Rosling A, Bever JD, Fransson P. 2016. Dominant mycorrhizal association of trees alters carbon and nutrient cycling by selecting for microbial groups with distinct enzyme function. New phytologist 214: 432–442.

Cheng L, Booker FL, Tu C, Burkey KO, Zhou L, Shew HD, Rufty TW, Hu S. 2012. Arbuscular mycorrhizal fungi increase organic carbon decomposition under elevated CO_2_. Science 337: 1084–1087.

Domeignoz-Horta LA, Shinfuku M, Junier P, Poirier S, Verrecchia E, Sebag D, DeAngelis KM. 2021. Direct evidence for the role of microbial community composition in the formation of soil organic matter composition and persistence. ISME Communications 1: 64.

Dumbrell AJ, Ashton PD, Aziz N, Feng G, Nelson M, Dytham C, Fitter AH, Helgason T. 2011. Distinct seasonal assemblages of arbuscular mycorrhizal fungi revealed by massively parallel pyrosequencing. New Phytologist 190: 794–804.

Dynarski KA, Bossio DA, Scow KM. 2020. Dynamic stability of soil carbon: Reassessing the “permanence” of soil carbon sequestration. Frontiers in Environmental Science 8: 514701.

Fossum C, Estera-Molina KY, Yuan M, Herman DJ, Chu-Jacoby I, Nico PS, Morrison KD, Pett-Ridge J, Firestone MK. 2022. Belowground allocation and dynamics of recently fixed plant carbon in a California annual grassland. Soil Biology and Biochemistry 165: 108519.

Frey SD. 2019. Mycorrhizal fungi as mediators of soil organic matter dynamics. *Annual Review of Ecology*, Evolution, and Systematics 50: 237–259.

Ghosal S, Fallon SJ, Leighton TJ, Wheeler KE, Kristo MJ, Hutcheon ID, Weber PK. 2008. Imaging and 3D elemental characterization of intact bacterial spores by high-resolution secondary ion mass spectrometry. Analytical chemistry 80: 5986–5992.

Godbold DL, Hoosbeek MR, Lukac M, Cotrufo MF, Janssens IA, Ceulemans R, Polle A, Velthorst EJ, Scarascia-Mugnozza G, De Angelis P et al. 2006. Mycorrhizal hyphal turnover as a dominant process for carbon input into soil organic matter. Plant and Soil 281: 15–24.

Graham EB, Knelman JE, Schindlbacher A, Siciliano S, Breulmann M, Yannarell A, Beman JM, Abell G, Philippot L, Prosser J et al. 2016. Microbes as engines of ecosystem function: When does community structure enhance predictions of ecosystem processes? Frontiers in Microbiology 7: 1–10.

Gui H, Purahong W, Hyde KD, Xu J, Mortimer PE. 2017. The arbuscular mycorrhizal fungus *Funneliformis mosseae* alters bacterial communities in subtropical forest soils during litter decomposition. Frontiers in Microbiology 8: 1120.

Habte M, Osorio NW. 2001. Arbuscular mycorrhizas: producing and applying arbuscular mycorrhizal inoculum. College of Tropical Agriculture and Human Resources, University of Hawaìi, Manoa. ISBN 1-929325-10-X

Halvorson JJ, Gonzalez JM. 2006. Bradford reactive soil protein in Appalachian soils: distribution and response to incubation, extraction reagent and tannins. Plant and Soil 286: 339–356.

Hao X, Zhu YG, Nybroe O, Nicolaisen MH. 2020. The composition and phosphorus cycling potential of bacterial communities associated with hyphae of *Penicillium* in soil are strongly affected by soil origin. Frontiers in microbiology, 10: 2951.

Hatton P-J., Kleber M, Zeller B, Moni C, Plante AF, Townsend K, Gelhaye L, Lajtha K, Derrien D. 2012. Transfer of litter-derived N to soil mineral–organic associations: evidence from decadal ^15^N tracer experiments. Organic Geochemistry 42: 1489–1501.

Hawkins HJ, Cargill RI, Van Nuland ME, Hagen SC, Field KJ, Sheldrake M, Soudzilovskaia NA, Kiers ET. 2023. Mycorrhizal mycelium as a global carbon pool. Current Biology 33: R560–R573.

Heckman K, Hicks Pries CE, Lawrence CR, Rasmussen C, Crow SE, Hoyt AM, von Fromm SF, Shi Z, Stoner S, McGrath C et al. 2022. Beyond bulk: Density fractions explain heterogeneity in global soil carbon abundance and persistence. Global Change Biology 28: 1178–1196.

Herman DJ, Firestone MK, Nuccio E, Hodge A. 2012. Interactions between an arbuscular mycorrhizal fungus and a soil microbial community mediating litter decomposition. FEMS Microbiology Ecology 80: 236–247.

Hestrin R, Kan M, Lafler M, Wollard J, Kimbrel JA, Ray P, Blazewicz S, Stuart R, Craven K, Firestone M et al. 2022. Plant-associated fungi support bacterial resilience following water limitation. The ISME Journal 16: 2752–2762.

Hicks Pries CE, Sulman BN, West C, O’Neill C, Poppleton E, Porras RC, Castanha C, Zhu B, Wiedemeier DB, Torn MS. 2018. Root litter decomposition slows with soil depth. Soil Biology and Biogeochemistry 125: 103–114.

Hodge A, Campbell C, Fitter A. 2001. An arbuscular mycorrhizal fungus accelerates decomposition and acquires nitrogen directly from organic material. Nature 413, 297–299.

Holátko J, Brtnický M, Kučerík J, Kotianová M, Elbl J, Kintl A, Kynický J, Benada O, Datta R, Jansa J. 2020. Glomalin – Truths, myths, and the future of this elusive soil glycoprotein. Soil Biology and Biochemistry 153: 108116.

Hooker JE, Piatti P, Cheshire MV, Watson CA. 2007. Polysaccharides and monosaccharides in the hyphosphere of the arbuscular mycorrhizal fungi *Glomus E3* and *Glomus tenue*. Soil Biology and Biochemistry 39: 680–683.

Horsch CCA, Antunes PM, Fahey C, Grandy AS, Kallenbach CM. 2023. Trait-based assembly of arbuscular mycorrhizal fungal communities determines soil carbon formation and retention. New Phytologist 239: 311–324.

Hoseinzade H, Ardakani MR, Shahdi A, Asadi Rahmani H, Noormohammadi G, Miransari M. 2016. Rice (*Oryza sativa L*.) nutrient management using mycorrhizal fungi and endophytic Herbaspirillum seropedicae. Journal of Integrative Agriculture 15: 1385–1394.

Hungate BA, Mau RL, Schwartz E, Caporaso, JG, Dijkstra P, van Gestel N, Koch BJ, Liu, CM, McHugh TA, Marks JC et al. 2015. Quantitative microbial ecology through stable isotope probing. Applied and Environmental Microbiology 81: 7570–7581.

Jakobsen I, Abbott LK, Robson AD. 1992. External hyphae of vesicular-arbuscular mycorrhizal fungi associated with *Trifolium subterraneum* L. 1. Spread of hyphae and phosphorus inflow into roots. New Phytologist 120: 371–380.

Jakobsen I, Rosendahl L. 1990. Carbon flow into soil and external hyphae from roots of mycorrhizal cucumber plants. New Phytologist 115: 77–83.

Jastrow JD, Amonette JE, Bailey VL. 2007. Mechanisms controlling soil carbon turnover and their potential application for enhancing carbon sequestration. Climate Change. 80: 5–23.

Jeffrey C. 1962. The origin and differentiation of the Archegoniate land-plants. Botaniska Notiser 115: 446–454.

Jiang F, Zhang L, Zhou J, George TS, Feng G. 2021. Arbuscular mycorrhizal fungi enhance mineralization of organic phosphorus by carrying bacteria along their extraradical hyphae. New Phytologist https://doi.org/10.1111/nph.17081.

Jones RT, Robeson MS, Lauber CL, Hamady M, Knight R, Fierer N. 2009. A comprehensive survey of soil acidobacterial diversity using pyrosequencing and clone library analyses. The ISME journal 3: 442–453.

Kaiser C, Kilburn MR, Clode PL, Fuchslueger L, Koranda M, Cliff JB, Solaiman ZM, Murphy DV. 2015. Exploring the transfer of recent plant photosynthates to soil microbes: mycorrhizal pathway vs direct root exudation. New Phytologist 205: 1537–1551.

Kakouridis A, Hagen JA, Kan MP, Mambelli S, Feldman LJ, Herman DJ, Weber PK, Pett-Ridge J, Firestone MK. 2022. Routes to roots: direct evidence of water transport by arbuscular mycorrhizal fungi to host plants. New Phytologist 236: 210–221.

Kallenbach CM, Frey SD, Grandy AS. 2016. Direct evidence for microbial-derived soil organic matter formation and its ecophysiological controls. Nature Communications 7: 13630.

Keiluweit M, Bougoure JJ, Nico PS, Pett-Ridge J, Weber PK, Kleber M. 2015. Mineral protection of soil carbon counteracted by root exudates. Nature Climate Change 5: 588–595.

Keiluweit M, Bougoure JJ, Zeglin LH, Myrold DD, Weber PK, Pett-Ridge J, Kleber M, Nico PS. 2012. Nano-scale investigation of the association of microbial nitrogen residues with iron (hydr)oxides in a forest soil O-horizon. Geochimica et Cosmochimica Acta 95: 213–226.

Kembel SW, Cowan PD, Helmus MR, Cornwell WK, Morlon H, Ackerly DD, Blomberg SP, Webb CO. 2010. Picante: R tools for integrating phylogenies and ecology. Bioinformatics 26: 1463–1464.

Kivlin SN, Fei S, Kalisz S, Averill C. 2020. Microbial ecology meets macroecology: developing a process-based understanding of the microbial role in global ecosystems. Bulletin Ecological Society of America 101: e01645.

Kleber M, Eusterhues K, Keiluweit M, Mikutta C, Mikutta R, Nico P. 2015. Mineral– organic associations: formation, properties, and relevance in soil environments. Advances in Agronomy 130: 1–140.

Kleber M, Sollins P, Sutton R. 2007. A conceptual model of organo-mineral interactions in soils: self-assembly of organic molecular fragments into zonal structures on mineral surfaces. Biogeochemistry 85: 9–24.

Koch, BJ, McHugh TA, Hayer M, Schwartz E, Blazewicz SJ, Dijkstra P, van Gestel N, Marks JC, Mau RL, Morrissey EM et al. 2018. Estimating taxon-specific population dynamics in intact microbial communities. Ecosphere 9: e02090

Kögel-Knabner I. 1997. ^13^C and ^15^N NMR spectroscopy as a tool in soil organic matter studies. Geoderma 80: 243–270.

Kögel-Knabner I. 2002. The macromolecular organic composition of plant and microbial residues as inputs to soil organic matter. Soil Biology and Biochemistry 34: 139–162.

Lang AK, Jevon FV, Vietorisz CR, Ayres MP, Hatala Matthes J. 2021. Fine roots and mycorrhizal fungi accelerate leaf litter decomposition in a northern hardwood forest regardless of dominant tree mycorrhizal associations. New Phytologist 230: 316–326.

Langley JA, Hungate BA. 2003. Mycorrhizal controls on belowground litter quality. Ecology 84: 2302–2312.

Leake J, Johnson D, Donnelly D, Muckle G, Boddy L, Read D. 2004. Networks of power and influence: the role of mycorrhizal mycelium in controlling plant communities and agroecosystem functioning. Canadian Journal of Botany 82: 1016–1045.

Lecomte J, St-Arnaud M, Hijri M. 2011. Isolation and identification of soil bacteria growing at the expense of arbuscular mycorrhizal fungi. FEMS Microbiology Letter 317: 43–51.

Lee J, Lee S, Young PW. 2008. Improved PCR primers for the detection and identification of arbuscular mycorrhizal fungi. FEMS Microbiology Ecology 65: 339–349.

Lehmann J, Hansel CM, Kaiser C, Kleber M, Maher K, Manzoni S, Nunan N, Reichstein M, Schimel JP, Manzoni S et al. 2020. Persistence of soil organic carbon caused by functional complexity. Nature Geoscience 13: 529–534.

Lei S, Xu X, Cheng Z, Xiong J, Ma R, Zhang L, Yang X, Zhu Y, Zhang B, & Tian B. 2019. Analysis of the community composition and bacterial diversity of the rhizosphere microbiome across different plant taxa. MicrobiologyOpen, 8(6): e00762.

Leifheit E, Verbruggen E, Rillig M. 2015. Arbuscular mycorrhizal fungi reduce decomposition of woody plant litter while increasing soil aggregation. Soil Biology and Biochemistry 13: 529– 534.

Li H, Bölscher T, Winnick M, Tfaily MM, Cardon ZG, Keiluweit M. 2021. Simple plant and microbial exudates destabilize mineral-associated organic matter via multiple pathways. Environmental Science & Technology 55: 3389–3398.

Love MI, Huber W, Anders S. 2014. Moderated estimation of fold change and dispersion for RNA-seq data with DESeq2. Genome Biology, 15: 550. doi:10.1186/s13059-014-0550-8.

Luthfiana N, Inamura N, Tantriani, Sato T, Oikawa A, Chen W, Tawaraya K. 2021. Metabolite profiling of the hyphal exudates of *Rhizophagus clarus* and *Rhizophagus irregularis* under phosphorus deficiency. Mycorrhiza 181: 1–10.

McGuire KL, Treseder KK. 2010. Microbial communities and their relevance for ecosystem models: Decomposition as a case study. Soil Biology Biochemistry 42: 529–535.

Miller RM, Jastrow JD. 2000. Mycorrhizal influence on soil structure. In: Kapulnik Y, Douds DD, eds. Arbuscular mycorrhizae: molecular biology and physiology. Dordrecht, the Netherlands: Kluwer Academic, 3–18.

Moni C, Rumpel C, Virto I, Chabbi A, Chenu C. 2010. Relative importance of sorption versus aggregation for organic matter storage in subsoil horizons of two contrasting soils. European Journal of Soil Science 61: 958–969.

Nacoon S, Jogloy S, Riddech N, Mongkolthanaruk W, Kuyper TW, Boonlue S. 2020. Interaction between Phosphate Solubilizing Bacteria and Arbuscular Mycorrhizal Fungi on Growth Promotion and Tuber Inulin Content of *Helianthus tuberosus* L. Scientific Reports 10: 4916.

Nuccio E, Blazewicz S, Lafler M, Campbell A, Kakouridis A, Kimbrel JA, Wollard J, Vyshenska D, Riley R, Tomatsu A et al. 2022. HT-SIP: A semi-automated Stable Isotope Probing pipeline identifies interactions in the hyphosphere of arbuscular mycorrhizal fungi. Microbiome 10: 199.

Nuccio EE, Hodge A, Pett-Ridge J, Herman DJ, Weber PK, Firestone MK. 2013. AMF alters soil bacterial community and N cycling. Environmental Microbiology 15: 1870–1881.

Oksanen J, Blanchet FG, Friendly M, Kindt R, Legendre P, McGlinn D, Minchin PR, O’hara RB, Simpson GL, Solymos P et al. 2019. Package ‘vegan’. Community ecology package, version 2.9.

Olsson PA, Wilhelmsson P. 2000. The growth of external AM fungal mycelium in sand dunes and in experimental systems. Plant Soil 226: 161–169.

Orwin KH, Kirschbaum MUF, St John MG, Dickie IA. 2011. Organic nutrient uptake by mycorrhizal fungi enhances ecosystem carbon storage: A model-based assessment. Ecology Letters 14: 493–502.

Parks D, Chuvochina M, Waite D et al. 2018. A standardized bacterial taxonomy based on genome phylogeny substantially revises the tree of life. Nature Biotechnology 36: 996–1004.

Parniske M. 2008. Arbuscular mycorrhiza: the mother of plant root endosymbioses. Nature Reviews Microbiology 6: 763–775.

Paterson E, Sim A, Davidson J, Daniell TJ. 2016. Arbuscular mycorrhizal hyphae promote priming of native soil organic matter mineralisation. Plant and Soil 408: 243–254.

Paul EA, Clark FE. 1989. Soil Microbiology and Biochemistry. Academic Press, San Diego.

Pett-Ridge J, Firestone MK. 2017. Using stable isotopes to explore root-microbe-mineral interactions in soil. Rhizosphere 3: 244–253.

Pett-Ridge J, Weber PK. 2012. NanoSIP: NanoSIMS applications for microbial biology. Microbial systems biology: methods and protocols: 375–408.

Pirozynski KA, Malloch DW. 1975. The origin of land plants: a matter of mycotrophism. BioSystems 6: 153–164.

Qin H, Niu L, Wu Q, Chen J, Li Y, Liang C, Xu Q, Fuhrmann JJ, Shen Y. 2017. Bamboo forest expansion increases soil organic carbon through its effect on soil arbuscular mycorrhizal fungal community and abundance. Plant Soil 420, 407–421.

Quiroga G, Erice G, Aroca R, Chaumont F, Ruiz-Lozano JM. 2019a. Contribution of the arbuscular mycorrhizal symbiosis to the regulation of radial root water transport in maize plants under water deficit. Environmental and Experimental Botany 167: 103821.

R. Core Team. 2017. R: A language and environment for statistical computing. Vienna, Austria.

Rodríguez-Caballero G, Caravaca F, Fernández-González AJ, Alguacil MM, Fernández-López M, Roldán A. 2017. Arbuscular mycorrhizal fungi inoculation mediated changes in rhizosphere bacterial community structure while promoting revegetation in a semiarid ecosystem. Science of the Total Environment 584: 838–848.

Rillig MC. 2004. Arbuscular mycorrhizae and terrestrial ecosystem processes. Ecological Letters 7: 740–754.

Rillig MC, Mardatin NF, Leifheit EF, Antunes PM. 2010. Mycelium of arbuscular mycorrhizal fungi increases soil water repellency and is sufficient to maintain water-stable soil aggregates. Soil Biological Biochemistry 42: 1189–1191.

Rillig MC, Mummey DL. 2006. Mycorrhizas and soil structure. New Phytologist 171: 41–53.

Rillig MC, Wright SF, Eviner VT. 2002. The role of arbuscular mycorrhizal fungi and glomalin in soil aggregation: comparing effects of five plant species. Plant Soil 238: 325–333

Rillig MC, Wright SF, Kimball BA, Pinter PJ, Wall GW, Ottman MJ, Leavitt SW. 2001b. Elevated carbon dioxide and irrigation effects on water stable aggregates in a sorghum field: a possible role for arbuscular mycorrhizal fungi. Global Change Biology 7: 333–337.

Rillig MC, Wright SF, Nichols KA, Schmidt WF, Torn MS. 2001a. Large contribution of arbuscular mycorrhizal fungi to soil carbon pools in tropical forest soils. Plant Soil 233: 167– 177.

Rorison B, Rorison IH. 1987. Root hairs and plant growth at low nitrogen availabilities. New Phytologist 107: 681–693.

Ruth B, Khalvati M, Schmidhalter U. 2011. Quantification of mycorrhizal water uptake via high-resolution on-line water content sensors. Plant and Soil 342: 459–468.

Sato T, Ezawa T, Cheng W, Tawaraya K. 2015. Release of acid phosphatase from extraradical hyphae of arbuscular mycorrhizal fungus *Rhizophagus clarus*. Journal of Soil Science and Plant Nutrition 61:269–274.

Schmidt MWI, Torn MS, Abiven S, Dittmar T, Guggenberger G, Janssens IA, Kleber M, Kögel-Knabner I, Lehman J, Manning DAC et al. 2011. Persistence of soil organic matter as an ecosystem property. Nature 478: 49–56.

Schüβler A, Schwarzott D, Walker C. 2001. A new fungal phylum, the Glomeromycota: phylogeny and evolution. Mycological Research 105:1413–1421.

See CR, Keller AB, Hobbie SE, Kennedy PG, Weber PK, Pett-Ridge J. 2022. Hyphae move matter and microbes to mineral microsites: integrating the hyphosphere into conceptual models of soil organic matter stabilization. Global Change Biology 8: 2527–2540.

Smith SE, Read DJ. 2008. Mycorrhizal symbiosis. Cambridge, UK: Academic Press.

Smits MM, Herrmann AM, Duane M, Duckworth OW, Bonneville S, Benning LG, Lundström U. 2009. The fungal–mineral interface: challenges and considerations of micro-analytical developments. Fungal Biology Reviews 23: 122–131.

Sokol N, Whalen ED, Jilling A, Kallenbach C, Pett-Ridge J, Georgiou K. 2022. The global distribution of mineral-associated soil organic matter, and its formation and fate under a changing climate. Functional Ecology 36: 1411–1429.

Sollins P, Kramer MG, Swanston C, Lajtha K, Filley T, Aufdenkampe AK, Wagai R, Bowden RD. 2009. Sequential density fractionation across soils of contrasting mineralogy: evidence for both microbial-and mineral-controlled soil organic matter stabilization. Biogeochemistry 96: 209–31.

Sollins P, Swanston C, Kleber M, Filley T, Kramer M, Crow S, Caldwell BA, Lajtha K, Bowden R. 2006. Organic C and N stabilization in a forest soil: evidence from sequential density fractionation. Soil Biology and Biochemistry 38: 3313–3324.

Soudzilovskaia NA, van Bodegom PM, Terrer C, van’t Zelfde M, McCallum I, McCormack ML, Fisher JB, Brundrett MC, de Sá NC, Tedersoo L. 2019. Global mycorrhizal plant distribution linked to terrestrial carbon stocks. Nature communications 10: 1– 10.

Soudzilovskaia NA, van der Heijden MG, Cornelissen JH, Makarov MI, Onipchenko VG, Maslov MN, Akhmetzhanova AA, van Bodegom PM. 2015. Quantitative assessment of the differential impacts of arbuscular and ectomycorrhiza on soil carbon cycling. New Phytologist 208: 280–93.

Spatafora JW, Chang Y, Benny GL, Lazarus K, Smith ME, Berbee ML, Bonito G, Corradi N, Grigoriev I, Gryganskyi A et al. 2016. A phylum-level phylogenetic classification of zygomycete fungi based on genome-scale data. Mycologia 108: 1028–46.

Strickland TC, Sollins P. 1987. Improved Method for Separating Light-and Heavy-Fraction Organic Material from Soil. Soil Science Society of America Journal 51: 1390–1393.

Sulman BN, Moore JA, Abramoff R, Averill C, Kivlin S, Georgiou K, Sridhar B, Hartman MD, Wang G, Wieder WR et al. 2018. Multiple models and experiments underscore large uncertainty in soil carbon dynamics. Biogeochemistry 141: 109–23.

Throckmorton HM, Bird JA, Monte N, Doane T, Firestone MK, Horwath WR. 2015. The soil matrix increases microbial C stabilization in temperate and tropical forest soils. Biogeochemistry. 122: 35–45.

Toljander JF, Artursson V, Paul LR, Jansson JK, Finlay RD. 2006. Attachment of different soil bacteria to arbuscular mycorrhizal fungal extraradical hyphae is determined by hyphal vitality and fungal species. FEMS Microbiology Letters 254: 34–40.

Toljander JF, Lindahl BD, Paul LR, Elfstrand M, Finlay RD. 2007. Influence of arbuscular mycorrhizal mycelial exudates on soil bacterial growth and community structure. FEMS Microbiology Ecology 61: 295–304.

Tome E, Tagliavini M, Scandellari F. 2015. Recently fixed carbon allocation in strawberry plants and concurrent inorganic nitrogen uptake through arbuscular mycorrhizal fungi. Journal of Plant Physiology 179: 83–89.

Torn MS, Trumbore SE, Chadwick OA, Vitousek PM, Hendricks DM. 1997. Mineral control of soil organic carbon storage and turnover. Nature 389: 170–173.

Treseder KK. 2016. Model behavior of arbuscular mycorrhizal fungi: Predicting soil carbon dynamics under climate change. Botany 94: 417–423.

Trumbore S. 2000. Age of soil organic matter and soil respiration: radiocarbon constraints on belowground C dynamics. Ecological applications: a publication of the Ecological Society of America 10: 399–411.

van der Heijden MGA, Martin FM, Selosse M-A, Sanders IR. 2015. Mycorrhizal ecology and evolution: the past, the present, and the future. New phytologist 205: 1406–1423.

Wagg C, Bender SF, Widmer F, van der Heijden MG. 2014. Soil biodiversity and soil community composition determine ecosystem multifunctionality. Proceedings of the National Academy of Sciences 111: 5266–5270.

Wang ZG, Bi YL, Jiang B, Zhakypbek Y, Peng SP, Liu WW, Liu H. 2016. Arbuscular mycorrhizal fungi enhance soil carbon sequestration in the coalfields, northwest China. Scientific reports 6: 1–11.

Wei L, Vosátka M, Cai B, Ding J, Lu C, Xu J, Yan W, Li Y, Liu C. 2019. The role of arbuscular mycorrhiza fungi in the decomposition of fresh residue and soil organic carbon: A mini-review. Soil Science Society of America Journal 83: 511–517.

Willis A, Rodrigues BF, Harris PJ. 2013. The ecology of arbuscular mycorrhizal fungi. Critical Reviews in Plant Sciences 32: 1–20.

Wilson GW, Rice CW, Rillig MC, Springer A, Hartnett DC. 2009. Soil aggregation and carbon sequestration are tightly correlated with the abundance of arbuscular mycorrhizal fungi: results from long-term field experiments. Ecological Letters 12: 452–461.

Wright SF, Franke-Snyder M, Morton JB, Upadhyaya A. 1996. Time-course study and partial characterization of a protein on hyphae of arbuscular mycorrhizal fungi during active colonization of roots. Plant Soil 181: 193–203.

Wu L, Wen C, Qin Y, Yin H, Tu Q, Van Nostrand JD, Yuan T, Yuan M, Deng Y, Zhou J. 2015. Phasing amplicon sequencing on Illumina Miseq for robust environmental microbial community analysis. BMC microbiology 15: 1–12.

Yuan MM, Kakouridis A, Starr E, Nguyen N, Shi S, Pett-Ridge J, Nuccio E, Zhou J, Firestone M. 2021. Fungal-bacterial co-occurrence patterns differ between AMF and non-mycorrhizal fungi across soil niches. MBio 12: 10–1128.

Zeglin LH, Wang B, Waythomas C, Rainey F, Talbot SL. 2016. Microbial structure and function after volcanic eruption. Environmental Microbiology 18: 146–158.

Zhang L, Fan J, Ding X, He XH, Zhang FS, Feng G. 2014. Hyphosphere interactions between an arbuscular mycorrhizal fungus and a phosphate solubilizing bacterium promote phytate mineralization in soil. Soil Biology and Biochemistry 74: 177–183.

Zhang HS, Zhou MX, Zai XM, Zhao FG, Qin P. 2020. Spatio-temporal dynamics of arbuscular mycorrhizal fungi and soil organic carbon in coastal saline soil of China. Scientific reports 1: 1–13.

