## Supplementary Information for "Arbuscular mycorrhiza convey significant plant carbon to a diverse hyphosphere microbial food web and mineral-associated organic matter"

The following Supporting Information is available for this article:

**Fig. S1.:** Atom% $^{13}C$ in shoots, roots, sand mix and soil mix in +AMF, -AMF, and $^{12}C$ microcosms.
**Fig. S2.:** Solid state NMR spectra of the soil mix (no-plant compartment) of +AMF and -AMF microcosms.
**Table S1.**: Data used in statistical analyses.
**Table S2**: Data used to calculate how much $^{13}C$ and total C was transported by AMF in +AMF microcosms.
**Table S3.**: Atom% $^{13}C$ of hyphae from NanoSIMS measurements for the 37 samples discussed in the main text.
**Table S4.:** ASVs that significantly increased or decreased in relative abundance in the soil mix (no-plant compartment) of +AMF microcosms.
**Method S1.**: Detailed $^{13}C$ calculations.

**Figure S1:** Atom% $^{13}C$ in shoots, roots, sand mix (plant compartment) and soil mix (no-plant compartment) in +AMF, -AMF, and $^{12}C$ microcosms. Different letters above bars represent statistically significant difference and corresponding p-values are indicated above the plots (one-way ANOVA & Fisher LSD test); a green line labeled "N" represents natural abundance levels of $^{13}C$.

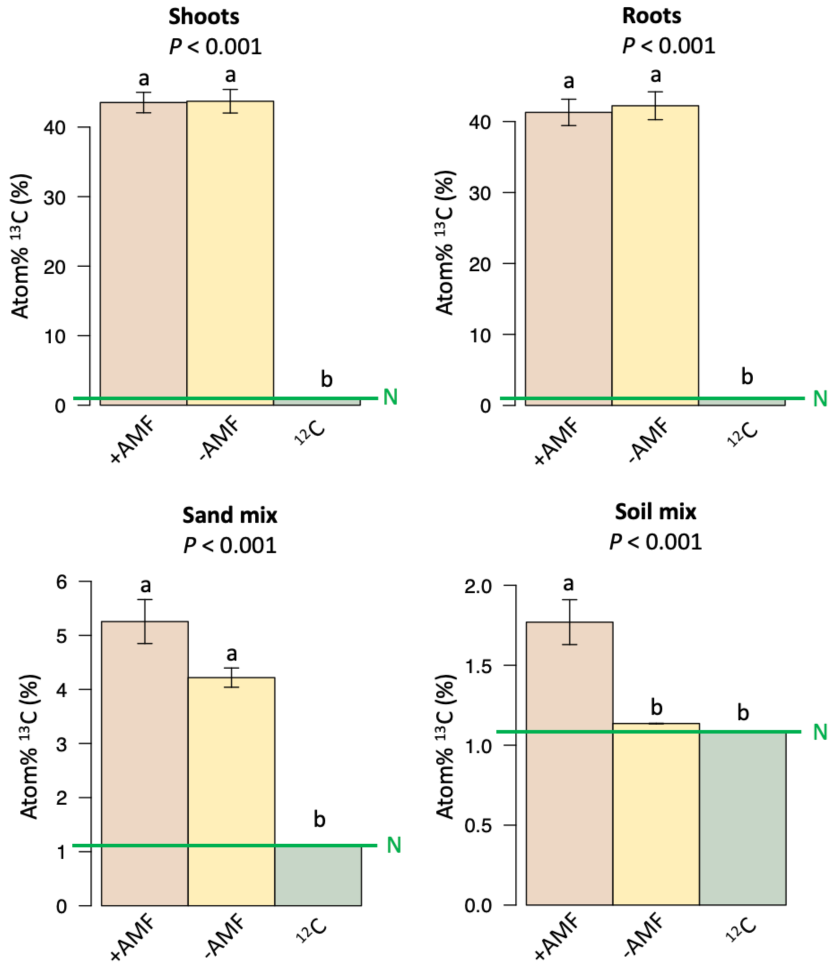

**Figure S2:** Solid state $^{13}$C NMR spectra of the soil mix (no-plant compartment) of +AMF microcosms (in orange, three replicates) and -AMF microcosms (in yellow, three replicates).

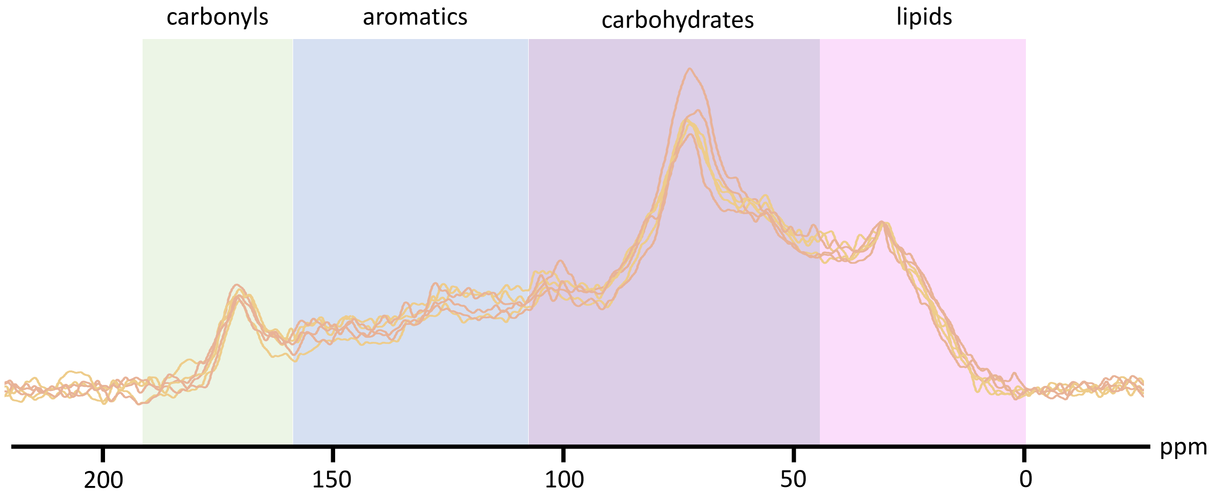

**Table S1:** Data used in statistical analyses. Values that appear in the main text are in **bold.** Means ± standard error are averages of twelve microcosms for +AMF treatment and six microcosms for -AMF and $^{12}$C treatments, each having three plants (one-way ANOVA & Fisher LSD test).

| | | "+AMF" | "-AMF" | "$^{12}$C" |
|---|---|---|---|---|
| plant compartment | mass of dry sand-clay mixture (g) | 745.1 | 745.1 | 745.1 |
| | gravimetric water content of sand-clay mixture at harvest (%) | 16.5 ±0.6 | 15.9 ±1.0 | 16.1 ±0.5 |
| | volume of plant compartment filled with sand-clay mixture (cm$^3$) | 612.5 | 612.5 | 612.5 |
| | above ground biomass at harvest (mg) | 559.2 ±64.5 | 535 ±49.7 | 769.2 ±76.6 |
| | below ground biomass at harvest (mg) | 805.3 ±80.5 | 801.9 ±75.2 | 814.5 ±88.8 |
| | above ground biomass %C at harvest (%) | 37.73 ±0.42 | 36.04 ±0.59 | 37.03 ±0.60 |
| | above ground biomass %N at harvest (%) | 0.89 ±0.02 | 0.77 ±0.12 | 0.81 ±0.02 |
| | above ground biomass C:N at harvest | 42.63 ±1.37 | 47.41 ±2.22 | 48.00 ±6.79 |
| | above ground biomass %P at harvest (%) | 0.29 ±0.05 | 0.12 ±0.012 | 0.25 ±0.012 |
| | **shoots average atom% $^{13}$C at harvest (%)** | **43.54 ±1.47** | **43.72 ±1.70** | **1.09 ±0.001** |
| | **roots average atom% $^{13}$C at harvest (%)** | **41.29 ±1.86** | **42.22 ±1.96** | **1.09 ±0.002** |
| | **sand-clay mix average atom%$^{13}$C at harvest (%)** | **5.26 ±0.41** | **4.22 ±0.18** | **1.1 ±0.003** |
| | total P added as Rorison's solution (µmol) | 10.11 | 20.22 | 10.11 |
| | total P added as bone meal (µmol) | 76.93 | 76.93 | 76.93 |
| | total N added as Rorison's solution (µmol) | 403.30 | 806.60 | 403.30 |
| | total N added as bone meal (µmol) | 111.38 | 111.38 | 111.38 |
| | total Fe added as Rorison's solution (µmol) | 6.82 | 13.62 | 6.82 |
| no-plant compartment | mass of dry soil-sand mixture (g) | 230.0 | 230.0 | 230.0 |
| | gravimetric water content of soil-sand mixture at harvest (%) | 11.6 ±0.3 | 11.1 ±1.1 | 9.9 ±0.4 |
| | volume of no-plant compartment filled with soil-sand mixture (cm$^3$) | 183.8 | 183.8 | 183.8 |
| | **soil-sand mix average atom%$^{13}$C at harvest (%)** | **1.77 ±0.14** | **1.14 ±0001** | **1.08 ±0.0003** |
| | **soil-sand mix average mg of $^{13}$C at harvest (mg)** | **66.37 ±3.19** | **33.83 ±1.59** | **39.63 ±0.71** |
| | total P added as Rorison's solution (µmol) | 101.1 | 101.1 | 101.1 |
| | total P added as bone meal (µmol) | 76.93 | 76.93 | 76.93 |
| | total N added as Rorison's solution (µmol) | 403.20 | 403.40 | 403.20 |
| | total N added as bone meal (µmol) | 111.38 | 111.38 | 111.38 |
| | total Fe added as Rorison's solution (µmol) | 6.82 | 6.82 | 6.82 |

**Table S2:** Data used to calculate how much C was transported by AMF in +AMF microcosms. Means ± standard error are averages of twelve microcosms for +AMF treatment and six microcosms for -AMF and $^{12}C$ treatments, each having three plants (one-way ANOVA & Fisher LSD test). Columns headers *A-J* represent letters used in the Methods **S1** calculations.

| | | | *A* | *B* | *C* | *D* | *E* | *F* | *G* | *H* | *I* | *J* |
|---|---|---|---|---|---|---|---|---|---|---|---|---|
| **Treatment** | | **Soil fraction** | **Average fraction weight in 20 g soil mix used for density fractonation (g)** | **Average %C (%)** | **Average atom% $^{13}C$ (%)** | **Average mg $^{13}C$ per microcosm (mg)** | **Average mg $^{13}C$ transported by AMF (mg)** | **Average total mg $^{13}C$ transported by AMF per microcosm (mg)** | **Average fraction weight in 20 g soil mix used for density fractonation (g)** | **Average %C (%)** | **Average mg C per microcosm (no-plant compartment) (mg)** | **Avg total mg C per microcosm (mg)** |
| **+AMF (no-plant compartment)** | $^{12}C$ | Light | 0.214±0.011 | 27.345±0.651 | 1.083±0.001 | 6.849±0.160 | **17.8±2.5** | **26.7±3.0** | 0.197±0.005 | 27.281±0.326 | 618.051±17.338 | 3621.706±115.694 |
| | $^{13}C$ | | 0.188±0.004 | 27.248±0.389 | 3.921±0.384 | 24.672±2.503 | | | | | | |
| | $^{12}C$ | Occluded | 0.227±0.017 | 35.167±0.460 | 1.080±0.0001 | 9.281±0.121 | **2.2±0.4** | | 0.208±0.008 | 34.567±0.350 | 826.843±32.885 | |
| | $^{13}C$ | | 0.198±0.007 | 34.267±0.460 | 1.364±0.031 | 11.432±0.334 | | | | | | |
| | $^{12}C$ | Heavy | 19.805±0.129 | 0.949±0.029 | 1.087±0.002 | 23.499±0.740 | **6.8±1.6** | | 19.800±0.069 | 0.956±0.048 | 2176.812±109.559 | |
| | $^{13}C$ | | 19.796±0.101 | 0.953±0.040 | 1.392±0.016 | 30.261±1.382 | | | | | | |

**Table S3:** Atom% $^{13}C$ of hyphae in +AMF microcosms from nanoscale secondary ion mass spectrometry (NanoSIMS) measurements for the 37 samples discussed in the main text.

| Microcosm # | Compartment | Sample content | Atom fraction | Atom fraction error | Atom% $^{13}C$ | Uncertainty |
|---|---|---|---|---|---|---|
| 1 | plant | hyphae attached to root 1 | 0.47996 | 0.00107 | 47.996 | 0.10720 |
| | | hyphae attached to root 2 | 0.42733 | 0.00118 | 42.733 | 0.11849 |
| | | hyphae attached to root 3 | 0.28995 | 0.00238 | 28.995 | 0.23831 |
| | | hyphae attached to root 4 | 0.40584 | 0.00540 | 40.584 | 0.54014 |
| | | hyphae attached to root 5 | 0.37201 | 0.02269 | 37.201 | 2.26901 |
| | | hyphae attached to root 6 | 0.46315 | 0.00048 | 46.315 | 0.04846 |
| 1 | no-plant | hyphae 1 | 0.61343 | 0.00092 | 61.343 | 0.09207 |
| | | hyphae 2 | 0.34955 | 0.00029 | 34.955 | 0.02934 |
| | | hyphae 3 | 0.53147 | 0.00030 | 53.147 | 0.02953 |
| | | hyphae 4 | 0.37293 | 0.00021 | 37.293 | 0.02109 |
| 2 | no-plant | hyphae 1 | 0.32225 | 0.00024 | 32.225 | 0.02411 |
| | | hyphae 2 | 0.31050 | 0.00028 | 31.050 | 0.02813 |
| | | hyphae 3 | 0.36547 | 0.00044 | 36.547 | 0.04379 |
| | | hyphae 4 | 0.43613 | 0.00109 | 43.613 | 0.10879 |
| | | hyphae 5 | 0.45055 | 0.00074 | 45.055 | 0.07397 |
| | | hyphae 6 | 0.33562 | 0.00025 | 33.562 | 0.02491 |
| | | hyphae 7 | 0.33285 | 0.00022 | 33.285 | 0.02249 |
| | | hyphae 8 | 0.46238 | 0.00088 | 46.238 | 0.08786 |
| | | hyphae 9 | 0.40317 | 0.00085 | 40.317 | 0.08525 |
| | | hyphae 10 | 0.39235 | 0.00234 | 39.235 | 0.23438 |
| | | hyphae 11 | 0.32361 | 0.00250 | 32.361 | 0.24989 |
| | | hyphae 12 | 0.29238 | 0.00139 | 29.238 | 0.13945 |
| | | hyphae 13 | 0.41712 | 0.00071 | 41.712 | 0.07133 |
| 3 | air gap | hyphae 1 | 0.34085 | 0.00114 | 34.085 | 0.11388 |
| | | hyphae 2 | 0.12437 | 0.00047 | 12.437 | 0.04669 |
| | | hyphae 3 | 0.34545 | 0.00277 | 34.545 | 0.27665 |
| | | hyphae 4 | 0.31999 | 0.00273 | 31.999 | 0.27282 |
| | | hyphae 5 | 0.35419 | 0.00065 | 35.419 | 0.06534 |
| | | hyphae 6 | 0.36759 | 0.00025 | 36.759 | 0.02533 |
| | | hyphae 7 | 0.32373 | 0.00036 | 32.373 | 0.03641 |
| | | hyphae 8 | 0.34591 | 0.00042 | 34.591 | 0.04184 |
| 4 | no-plant | hyphae 1 | 0.23931 | 0.00078 | 23.931 | 0.07849 |
| | | hyphae 2 | 0.26367 | 0.00026 | 26.367 | 0.02621 |
| | | hyphae 3 | 0.25446 | 0.00041 | 25.446 | 0.04102 |
| | | hyphae 4 | 0.26520 | 0.00023 | 26.520 | 0.02342 |
| | | hyphae 5 | 0.23993 | 0.00076 | 23.993 | 0.07610 |
| 6 | no-plant | hyphae 1 | 0.33941 | 0.00057 | 33.941 | 0.05742 |

**Table S4:** ASVs that significantly **(a)** increased and **(b)** decreased in relative abundance (by DESeq analysis of 16S rRNA reads, $P < 0.05$ adjusted for multiple comparisons) in the soil mix (no-plant compartment) of +AMF microcosms compared to ASVs in the soil mix of -AMF microcosms.

**(a)**

| ASVs that increased in relative abundance in the presence of AMF | | | | | | | |
|---|---|---|---|---|---|---|---|
| log2FoldChange | ASV | Domain | Phylum | Class | Order | Family | Genus |
| 5.299132181 | bdbdb4a39e3433bcdab5de2db131d9d6 | Bacteria | Actinobacteria | Actinobacteria | Micrococcales | Micrococcaceae | |
| 6.823631441 | 93b85a70b948a8c9b82f618b0ab9f575 | Bacteria | Bacteroidetes | Bacteroidia | Sphingobacteriales | env.OPS 17 | |
| 4.707805791 | 250aa457e3eae7fea4e7936c3ff524aa | Bacteria | Chloroflexi | Anaerolineae | SBR1031 | A4b | |
| 7.175240394 | 79029fa983ec4eda60117a273a52cfc0 | Bacteria | Proteobacteria | Alphaproteobacteria | Caulobacterales | Caulobacteraceae | Caulobacter |
| 6.799645131 | 971eb60b1020447dc394509ffc8c25d4 | Bacteria | Proteobacteria | Alphaproteobacteria | Dongiales | Dongiaceae | Dongia |
| 3.505753586 | 2743823e7602e70cfa467627e69d2774 | Bacteria | Proteobacteria | Alphaproteobacteria | Rhizobiales | Rhizobiaceae | Allorhizobium-Neorhizobium-Pararhizobium-Rhizobium |
| 6.514034761 | a9164d01308501b576afa06256a15b0f | Bacteria | Proteobacteria | Alphaproteobacteria | Rhizobiales | Rhizobiaceae | Allorhizobium-Neorhizobium-Pararhizobium-Rhizobium |
| 6.781343603 | adbda4915419b8f4cd4a001a04cdfc02 | Bacteria | Proteobacteria | Alphaproteobacteria | Rhizobiales | Xanthobacteraceae | Tardiphaga |
| 5.433349515 | 146011b4b3b1ac5bd7d8224c5d93209d | Bacteria | Proteobacteria | Alphaproteobacteria | Sphingomonadales | Sphingomonadaceae | Sphingomonas |
| 7.341846244 | 396c2d79acbe8883549b9049dddce9fb | Bacteria | Proteobacteria | Alphaproteobacteria | Sphingomonadales | Sphingomonadaceae | Sphingomonas |
| 6.831396962 | ef20b3cbed3170b3a8faf33cf0b1db59 | Bacteria | Proteobacteria | Alphaproteobacteria | Sphingomonadales | Sphingomonadaceae | Sphingomonas |
| 7.035481394 | e8a499c569880b49145fc0bb9a2f71c5 | Bacteria | Proteobacteria | Deltaproteobacteria | Myxococcales | Myxococcaceae | Myxococcus |
| 6.567182906 | 78d2353d11fe9e4a8a5976115cf6ae57 | Bacteria | Proteobacteria | Deltaproteobacteria | Myxococcales | | |
| 7.861976969 | 3a09c6c2ffa5e76cdc6b6d1324df0a0b | Bacteria | Proteobacteria | Gammaproteobacteria | Betaproteobacteriales | Burkholderiaceae | Herbaspirillum |
| 9.017622164 | 07b9db0375df4a5e63c8a479145d4478 | Bacteria | Proteobacteria | Gammaproteobacteria | Betaproteobacteriales | Burkholderiaceae | |
| 29.97971268 | 58c1e669ce1aaeedf8bd611fc28d6cba | Bacteria | Proteobacteria | Gammaproteobacteria | Betaproteobacteriales | Burkholderiaceae | |
| 7.382460074 | f7e814485a8a76f48d6cff53ff83e798 | Bacteria | Proteobacteria | Gammaproteobacteria | Betaproteobacteriales | Burkholderiaceae | |
| 7.617530768 | 9764767233c1b1ea3dc4932b94be8512 | Bacteria | Verrucomicrobia | Verrucomicrobiae | Chthoniobacterales | Chthoniobacteraceae | Chthoniobacter |
| 6.513961545 | 1df9396534c5a37ad221a45747e73a89 | Bacteria | Verrucomicrobia | Verrucomicrobiae | UA11 | | |

**(b)**

| ASVs that decreased in relative abundance in the presence of AMF | | | | | | | |
|---|---|---|---|---|---|---|---|
| log2FoldChange | ASV | Domain | Phylum | Class | Order | Family | Genus |
| -6.7873212 | 0be31311933b0d26d811dd5df11c8116 | Bacteria | Acidobacteria | Subgroup 6 | uncultured Anaeromyxo | uncultured Anaeromyx | uncultured Anaeromyxobacter sp. |
| -5.12202 | 25da1a2223a07f6386538f347b59b16b | Bacteria | Bacteroidetes | Bacteroidia | Cytophagales | Cytophagaceae | Sporocytophaga |
| -5.4279578 | 391cb53f747f95a40325699d209bcec8 | Bacteria | Bacteroidetes | Bacteroidia | Cytophagales | Microscillaceae | |
| -6.1828696 | 8b5e82b40cc5dd728caeb1da7aa941f7 | Bacteria | Cyanobacteria | Melainabacteria | Vampirovibrionales | | |
| -7.4569919 | 732b608a5a32f5265c7f84a8e337f8c4 | Bacteria | Fibrobacteres | Fibrobacteria | Fibrobacterales | Fibrobacteraceae | possible genus 04 |
| -5.5685953 | d02d26187e182c606b9dbb0cf06941c2 | Bacteria | Gemmatimonade | S0134 terrestrial group | | | |
| -3.7071829 | 229de90b8ae8b2f8587fe7f9cb223083 | Bacteria | Patescibacteria | Saccharimonadia | Saccharimonadales | | |
| -6.8634864 | 4e6aa4bfcc393ce0631afd7ab8754d21 | Bacteria | Planctomycetes | vadinHA49 | | | |
| -5.7545578 | 52f7dd3bd77e7b9830c88dda40dc695d | Bacteria | Proteobacteria | Alphaproteobacteria | Micropepsales | Micropepsaceae | uncultured Hyphomicrobiaceae bacterium |
| -7.4856253 | 23d450899c1481b4f8601c28fc70dcb3 | Bacteria | Proteobacteria | Alphaproteobacteria | Rhizobiales | Xanthobacteraceae | Pseudolabrys |
| -7.4550547 | 5901d848a9dc04b3c0236b4011412a50 | Bacteria | Proteobacteria | Deltaproteobacteria | Myxococcales | BIrii41 | |
| -6.7802369 | ed4897364210960229ae8b3074c5a6ed | Bacteria | Proteobacteria | Deltaproteobacteria | Myxococcales | Haliangiaceae | Haliangium |
| -7.9449615 | 5f8db894a571d35c9d2fa4f8df8581f9 | Bacteria | Proteobacteria | Gammaproteobacteria | Betaproteobacteriales | Burkholderiaceae | Massilia |
| -5.6810353 | caa187b77f27c28a11a913a519da60b9 | Bacteria | Proteobacteria | Gammaproteobacteria | Betaproteobacteriales | Burkholderiaceae | Massilia |
| -6.9822615 | d4cb29080e56a06ef36f99b209f2b225 | Bacteria | Proteobacteria | Gammaproteobacteria | Betaproteobacteriales | Nitrosomonadaceae | MND1 |
| -7.242162 | df71de0e92459aa808364311135bb8fc | Bacteria | Proteobacteria | Gammaproteobacteria | Betaproteobacteriales | Burkholderiaceae | Variovorax |
| -5.9222471 | c35f665bd95ab81a5ca34ee89524dba5 | Bacteria | Proteobacteria | Gammaproteobacteria | Betaproteobacteriales | Burkholderiaceae | |

**Method S1**: $^{13}C$ calculations

These calculations determine 1) how much AMF-transported C ($^{13}C$ or $^{12}C$ + $^{13}C$ combined) that crossed the air gap remained after losses as respiration, DOC, etc., and 2) what percent of total soil C the AMF-transported C represents.

Each no-plant compartment contained 230 g of soil mix ($m_{soil\_mix}$), which was made up of 133 g of sand ($m_{sand}$) and 97 g of soil ($m_{soil}$). A 20 g soil mix was sampled ($m_{sample}$) for density fractionation and subsequent C and $^{13}C$ analysis from each microcosm. All weights are dry weights. The averages and standard errors of the raw measurements and calculated values are reported in Table **S2**.

Letters used in the calculations below represent columns in Table **S2**. For the 20 g subsample from each microcosm, we measured the weight of each density fraction, $\boldsymbol{A}$. For each density fraction, we measured the %C, $\boldsymbol{B}$ and atom% $^{13}C$, $\boldsymbol{C}$.

First, we calculated the amount of $^{13}C$ in each fraction of each microcosm's no-plant compartment, $\boldsymbol{D}$, in mg, by scaling up the amount of $^{13}C$ in the samples we analyzed:

$$\boldsymbol{D} = \boldsymbol{A} \times \frac{m_{soil\_mix}}{m_{sample}} \times \frac{\boldsymbol{B}}{100} \times \frac{\boldsymbol{C}}{100} \times 1000$$

The amount of $^{13}C$ contributed by AMF to each fraction, $\boldsymbol{E}$, in mg, was subsequently calculated by subtracting the amount of $^{13}C$ in the $^{12}C$ microcosms of that in the $^{13}C$ microcosm for each fraction:

$$\boldsymbol{E} = \boldsymbol{D}_{13C} - \boldsymbol{D}_{12C}$$

Adding the fractions up, we got the total amount of $^{13}C$ in each microcosm that was contributed by AMF, $\boldsymbol{F}$, in mg:

$$\boldsymbol{F} = \boldsymbol{E}_{light} + \boldsymbol{E}_{occluded} + \boldsymbol{E}_{heavy}$$

Similarly, we calculated the total amount of C in each microcosm, $\boldsymbol{J}$, in mg, by considering the average weight of each fraction across $^{12}C$ and $^{13}C$ microcosms, $\boldsymbol{G}$, in g, average %C, $\boldsymbol{H}$, and sum up the amount of C in each the three fractions $\boldsymbol{I}$, in mg:

$$\boldsymbol{I} = \boldsymbol{G} \times \frac{\boldsymbol{H}}{100} \times \frac{m_{soil\_mix}}{m_{sample}} \times 1000$$

$$\boldsymbol{J} = \boldsymbol{I}_{light} + \boldsymbol{I}_{occluded} + \boldsymbol{I}_{heavy}$$

On average, there was $\boldsymbol{J}$ = 3619.3 mg of total C in no-plant compartment of the microcosms.

On average, after 6 weeks of labeling, the amount of $^{13}C$ AMF transported that remained in the soil was $\boldsymbol{F}$ = 26.7 mg. NanoSIMS data revealed that on average hyphae were $\boldsymbol{K}$ = 35.9% $^{13}C$, so the amount of total C ($^{12}C$ + $^{13}C$) AMF transported that remained in soil, $\boldsymbol{L}$, was

$$\boldsymbol{L} = \frac{\boldsymbol{F}}{\boldsymbol{K}} = 74.4 \text{ mg CA}$$

This is equivalent to

$$\frac{\boldsymbol{L}}{m_{\text{soil}}} = 0.77 \text{ mg C per g soil}$$

or

$$\frac{\boldsymbol{L}}{\boldsymbol{J}} = 2.05\% \text{ of total C in soil mix}$$

For these calculations, we used the following assumptions:

**A1**: Subsamples analyzed are representative of the C content, proportions of light, occluded, and heavy fractions, and $^{13}C$ enrichment across the entire soil mix in the no-plant compartment for each microcosm.

Justification: After preserving small amount of soil mix for molecular analysis and microscopy, we homogenized the entire content in the no-plant compartment and then took the subsample for C analysis.

**A2**: The enrichment level of AMF hyphae captured by NanoSIMS is representative, as least not overestimating, the level of $^{13}C$ enrichment in the C that AMF transported and remained in soil.

Justification: The $^{13}C$ enrichment in AMF hyphae was estimated by 37 images of hyphae sampled across different microcosms and locations. The material "piped" by AMF hyphae is likely more $^{13}C$ enriched than the hyphae themselves. In addition, soil respiration favors $^{12}C$ over $^{13}C$, so the back-calculated total C transported via AMF and remained in soil should be a conservative estimation.
